# Pancreas resident macrophage-induced fibrosis has divergent roles in pancreas inflammatory injury and PDAC

**DOI:** 10.1101/2022.02.09.479745

**Authors:** John M. Baer, Chong Zuo, Liang-I Kang, Angela Alarcon de la Lastra, Nicholas C. Borcherding, Brett L. Knolhoff, Savannah J. Bogner, Yu Zhu, Mark A. Lewis, Nan Zhang, Ki-Wook Kim, Ryan C. Fields, Jason C. Mills, Li Ding, Gwendalyn J. Randolph, David G. DeNardo

## Abstract

Tissue-resident macrophages (TRMs) are long-lived cells that maintain locally and can be phenotypically distinct from monocyte-derived macrophages (MDMs). However, whether TRMs and MDMs have functional distinction under differing pathologies is not understood. Here, we show a significant portion of macrophages that accumulated during pancreatitis and pancreatic cancer were expanded from TRMs. We further established that pancreas TRMs have a distinct extracellular matrix remodeling phenotype that was critical for maintaining tissue homeostasis during inflammation. Loss of TRMs led to exacerbation of severe pancreatitis and animal death, due to impaired acinar cell survival and recovery. In pancreatitis, TRMs elicited protective effects by triggering the accumulation and activation of fibroblasts, which was necessary for initiating fibrosis as a wound healing response. The same TRM-driven fibrosis, however, drove pancreas cancer pathogenesis and progression. Together, these findings indicate that TRMs play divergent roles in the pathogenesis of pancreatitis and cancer through regulation of stromagenesis.

## Introduction

Tissue-resident macrophages (TRMs) are established through waves of embryonic and adult hematopoiesis. At birth, all macrophages are derived from embryonic progenitors, but some tissue macrophages are gradually replaced with monocyte-derived cells over time, to varying extents determined at a tissue-specific level (Bain et al., 2016; Epelman et al., 2014a; Ginhoux et al., 2010; Ginhoux and Guilliams, 2016; Hashimoto et al., 2013; Hoeffel et al., 2015; Liu et al., 2019; Schulz et al., 2012; Yona et al., 2013). “Closed” TRM populations, such as brain microglia and liver Kupffer cells, undergo little to no replacement from monocytes, whereas other more “open” TRM populations, such as lung alveolar macrophages and pancreas macrophages, undergo gradual replacement, the extent, and rate of which is determined at a tissue-specific level (Calderon et al., 2015; Ginhoux and Guilliams, 2016; Liu *et al*., 2019). Despite heterogeneity in ontogeny between TRM populations within different tissues, they are defined by their ability to maintain their numbers locally through longevity and self-renewal. However, there are also populations of short-lived monocyte-derived macrophages (MDMs) that rely on continual replenishment from monocytes coming from hematopoietic stem cells in the bone marrow (Bain et al., 2014; Kim et al., 2016). Further, it is understood that both developmental origin and the local tissue microenvironment offer cues that contribute to the phenotype and function of macrophages (Lavin et al., 2014; Loyher et al., 2018; Zhu et al., 2017). TRMs have been implicated in a wide variety of roles in maintaining tissue homeostasis, extracellular matrix (ECM) remodeling, and inflammation (Ginhoux and Jung, 2014). Additionally, across the fat, dermis, heart, lung, and mesenteric membranes, a dogma has emerged that interstitial TRMs can be subdivided into at least two major subpopulations, Lyve1^Hi^CX3CR1^Lo^MHCII^Lo^ or Lyve1^Lo^CX3CR1^Hi^MHCII^Hi^ subsets, with multiple reports now implicating Lyve1^Hi^CX3CR1^Lo^MHCII^Lo^ TRMs as being involved in ECM remodeling (Chakarov et al., 2019; Dick et al., 2022; Lim et al., 2018; Zhang et al., 2021). However, it is not fully understood how TRM subsets and MDMs might be uniquely poised to respond to tissue damage or pathologic conditions and how their responses might differ.

Several studies have now investigated the potential roles of TRMs and MDMs under tumor conditions (Casanova-Acebes et al., 2021; Loyher *et al*., 2018; Zhu *et al*., 2017). TRMs have been identified in playing various tumor-promoting roles, such as driving regulatory T cell responses, contributing to fibrosis and ECM deposition, and supporting tumor cell growth (Casanova-Acebes *et al*., 2021; Loyher *et al*., 2018; Zhu *et al*., 2017). Our previous study has shown in pancreatic ductal adenocarcinoma (PDAC) that macrophages have divergent roles based on developmental origin, with embryonic-derived TRMs being particularly adept at driving fibrosis (Zhu *et al*., 2017). It is not understood, however, how these roles might be altered in different models of tissue injury. Inflammation and tissue damage during pancreatitis is not only a risk factor for PDAC, but also involves significant accumulation of desmoplastic stroma rich in myeloid cells (Apte et al., 2011; Klöppel et al., 2003). Thus far, macrophages have been implicated in contributing to inflammation during pancreatitis, but specific subsets of TRMs or MDMs have not been well characterized (Liou et al., 2013; Saeki et al., 2012; Xue et al., 2014). Studies have focused on how infiltrating macrophages can drive inflammation, but the roles of TRMs during pancreatitis are not well established. Additionally, the pancreas is a unique organ that rapidly responds with significant desmoplasia during tissue injury. While this fibrotic response is likely critical for preserving organ function, it has been shown to be a significant feature driving pancreas cancer progression and resistance to therapy (Jiang et al., 2017; Omary et al., 2007; Zimmermann, 2002).

Here, we show that both TRM and MDM subsets of macrophages in the pancreas increase drastically during pancreatitis and display distinct transcriptional phenotypes. We identify a Lyve1^Hi^ TRM subset expressing extracellular matrix remodeling and collagen genes, and further show that depleting TRMs negatively impacts mouse survival from severe acute pancreatitis by attenuating the fibrotic response. TRMs were shown to trigger the accumulation of fibroblasts, as well as affect their activation and stimulate the production of collagen and ECM proteins. While TRM-driven fibrosis played a protective role in pancreatitis, this was co-opted by tumors to support their growth.

## Results

### Human and mouse pancreatitis and PDAC display features of ADM and macrophage accumulation

Both pancreatitis and pancreatic ductal adenocarcinoma (PDAC) are characterized by desmoplastic fibrotic stromal responses and significant immune infiltration. To profile changes in the pancreas microenvironment during these pathologies, we first analyzed human tissue samples of adjacent normal pancreas, pancreatitis, and PDAC patients. Histological analysis of showed marked loss of acinar cell area in pancreatitis, replaced with fibrotic stroma (**Fig. 1a**), compared to normal pancreas tissue. To quantify macrophage numbers across disease states in human pancreas tissue, we co-immunostained for the macrophage marker CD163 in combination with cytokeratin 17/19 (CK17/19) to mark acinar areas, as well as cells undergoing acinar-to-ductal metaplasia (ADM) and tumor cells. CD163-expressing macrophages were significantly increased in in pancreatitis and PDAC samples, when compared to areas of normal pancreas (**Fig. 1a-b**). Additionally, to further examine the stromal alterations across disease states, we utilized trichrome and podoplanin staining to mark collagen and fibroblasts respectively **(Fig. 1a-b**). As with macrophage numbers, pancreatitis and PDAC samples showed significant increases in both collagen and fibroblasts compared to normal pancreas tissue.

**Figure 1:**
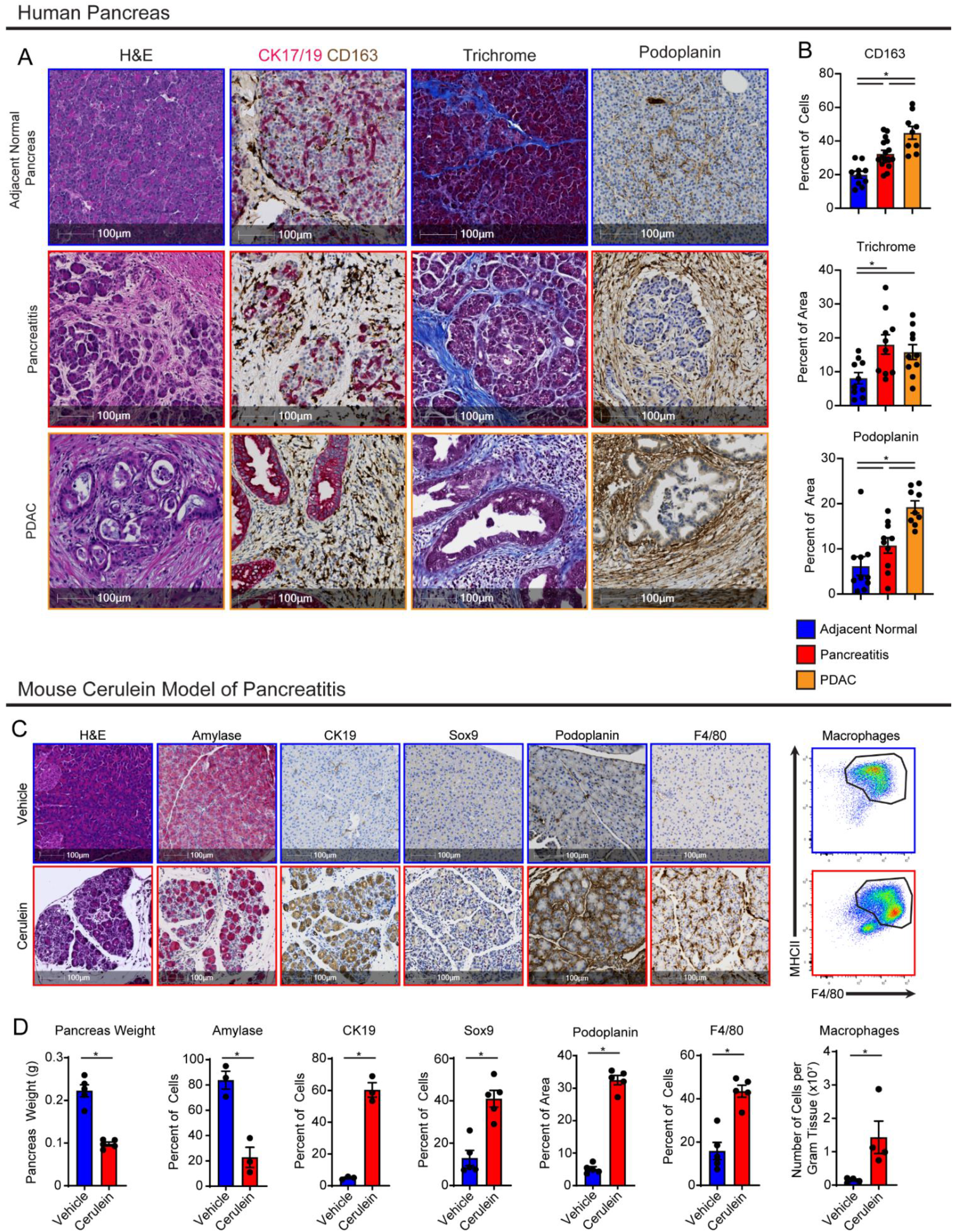
Human and mouse pancreatitis and PDAC display features of ADM and macrophage accumulation. **A**. Representative immunohistochemistry (IHC) images of H&E, CK17/19, CD163, trichrome, and podoplanin stains on human adjacent normal pancreas, pancreatitis, and PDAC tissue. **B**. Quantification of human adjacent normal pancreas, pancreatitis, and PDAC tissue positive for CD163, trichrome collagen (blue staining), or podoplanin staining, represented as percentage of cells or percentage of tissue area; n = 9-16 samples/group. **C**. Representative IHC images of pancreas tissue of mice treated with vehicle or cerulein stained for H&E, Amylase, CK19, Sox9, podoplanin, and F4/80, and flow cytometry plot of macrophages from pancreas tissue of vehicle or cerulein treated mice. **D**. Quantification of pancreas weight, Amylase, CK19, Sox9, podoplanin, and F4/80 IHC stains, and flow cytometry analysis of pancreas macrophages; n = 3-5 mice/group. Data are presented as mean ± SEM. n.s., not significant; *p <0.05. For comparisons between two groups, Student’s two-tailed t-test was used.

To better study pancreas injury and the impact of tissue resident macrophages (TRMs), we utilized the common cerulein-induced model of pancreatitis (Lerch and Gorelick, 2013). Serial injection of cerulein in mice is known to cause overproduction of digestive enzymes leading to autodigestion of pancreas tissue, acinar cells undergoing ADM, and infiltration of inflammatory immune cells. Compared to vehicle-treated mice, cerulein-injected mice showed widespread ADM by histology, and increased proportion of epithelial cells expressing common ADM markers Cytokeratin-19 (CK19) and Sox9 (**Fig. 1c-d**). Acinar cells undergoing ADM also have been shown to re-enter cell cycle, which we confirmed by increase in cells expressing Ki-67 (**Supp. Fig. 1c**). Next, to determine if macrophages accumulate in the pancreas during inflammatory injury caused by cerulein treatment, we stained F4/80 by IHC, which showed drastic accumulation of macrophages compared to vehicle-treated mice (**Fig. 1c-d**). Similarly, flow cytometry quantification of F4/80^+^MHCII^HI/Low^ macrophages showed a 10-fold increase in absolute number in cerulein-treated mice compared to vehicle-treated mice (**Fig. 1c-d, Supp. Fig. 1a**). Consistent with cerulein treatment causing widespread inflammation in the pancreas, we also observed increases in eosinophils, granulocytes, and monocytes in the pancreas with cerulein treatment (**Supp. Fig. 1b**). Collectively, our data show that cerulein-treated mice have similar accumulation of macrophages and ADM as observed in pancreatitis or PDAC patients.

### Cerulein treatment causes expansion of embryonic-derived, tissue-resident macrophages

We next sought to determine if the increase in macrophage number during pancreatitis was from monocytes or the local expansion of tissue-resident macrophages. First, to mark embryonic-derived macrophages, we used Flt3-Cre mice crossed to R26-LSL-eYFP mice (Flt3-YFP mice), which have previously been shown to mark all haematopoietically-derived cells with YFP, while embryonic-derived macrophages lack any YFP labeling (Boyer et al., 2011). In these mice, we observed that while both YFP-positive (Flt3-YFP^+^) macrophages and YFP-negative (Flt3-YFP^(-)^) macrophages were increased in cerulein-treated mice, Flt3-YFP^(-)^ macrophages was more markedly increased, showing 13-folds in absolute numbers. (**Fig. 2a-c**). Interestingly, Flt3-YFP^(-)^ macrophages show the largest increase in proliferation as marked by Ki-67 during cerulein treatment (**Fig. 2d**), consistent with TRMs increasing their numbers through *in situ* proliferation.

**Figure 2:**
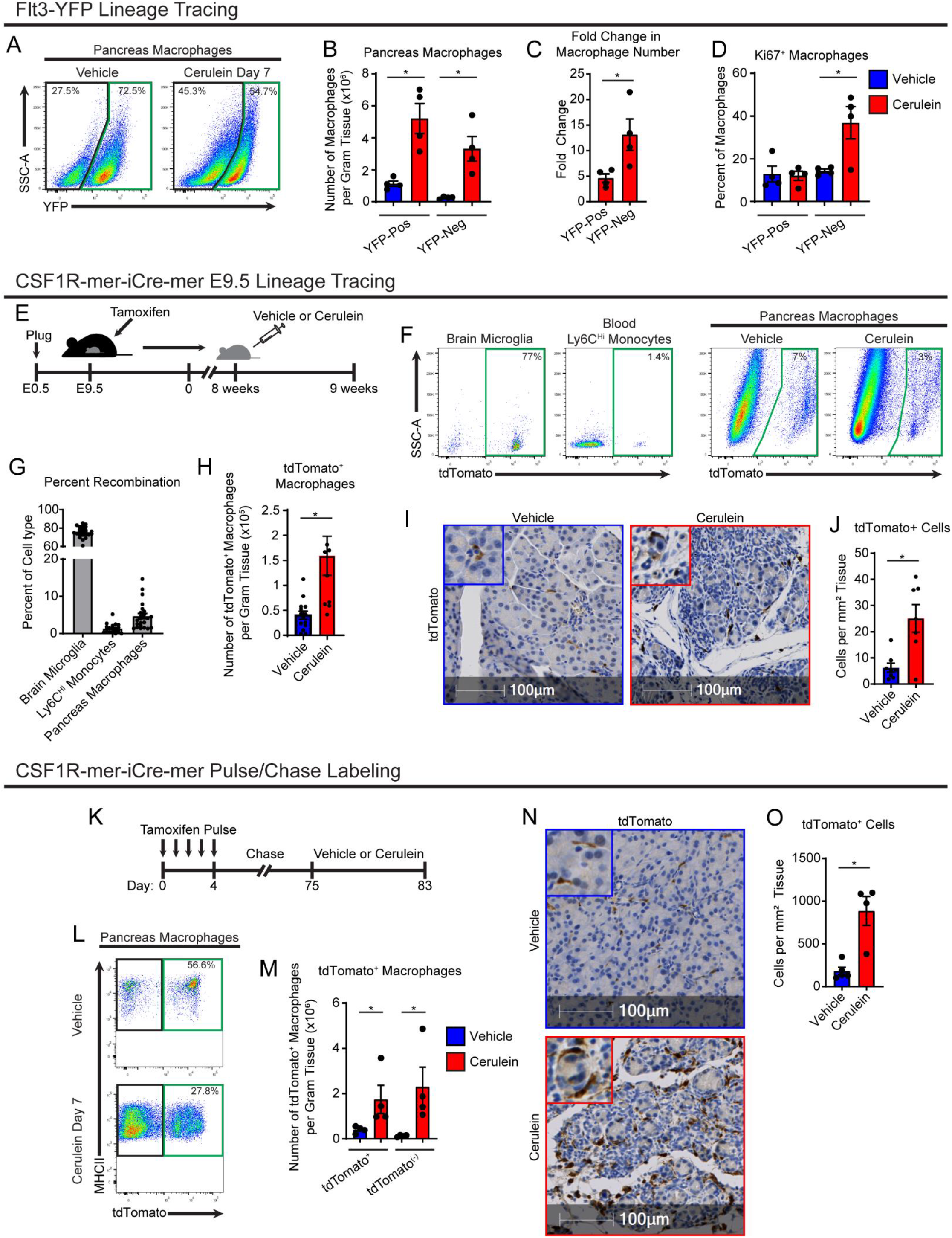
Cerulein treatment causes expansion of embryonic-derived, tissue-resident macrophages. **A**. Lineage tracing analysis of Flt3-Cre;LSL-YFP mice, displaying flow cytometry plots pre-gated on pancreas macrophages. **B**. Density of Flt3-YFP positive and negative macrophages in pancreas of vehicle or cerulein treated mice; n = 4 mice/group. **C**. Fold-change in density of Flt3-YFP positive and negative macrophages between vehicle and cerulein groups; n = 4 mice/group. **D**. Percentage of Flt3-YFP positive and negative macrophages positive for Ki-67 between vehicle and cerulein groups; n = 4 mice/group. **E**. Treatment scheme for embryonic pulse of tamoxifen in CSF1R-mer-iCre-mer;LSL-tdTomato mice. **F**. Flow cytometry plots of brain microglia, blood Ly6C^+^ monocytes, and vehicle or cerulein treated pancreas macrophages. **G**. Percentage of indicated cell type labeled by embryonic tamoxifen pulse; n = 24 mice/cell type analyzed. **H**. Density of pancreas macrophages labeled by embryonic tamoxifen pulse; n = 8-14 mice/group. **I**. Representative images of IHC stain for tdTomato on pancreas tissue of CSF1R-mer-iCre-mer;LSL-tdTomato mice treated with vehicle or cerulein. **J**. Quantification of pancreas macrophages labeled by tdTomato from embryonic tamoxifen pulse; n = 7-8 mice/group. **K**. Treatment scheme for tamoxifen pulse/chase labeling in CSF1R-mer-iCre-mer;LSL-tdTomato mice. **L**. Flow cytometry plots of tdTomato labeling in pancreas macrophages; n = 4 mice/group. **M**. Density of tdTomato^+^ and tdTomato^(-)^ pancreas macrophages in vehicle and cerulein treated groups; n = 4 mice/group. **N**. Representative images of IHC stain for tdTomato on pancreas tissue of CSF1R-mer-iCre-mer;LSL-tdTomato tamoxifen pulse/chase mice; n = 4 mice/group. **J**. Quantification of pancreas macrophages labeled by tdTomato from tamoxifen pulse/chase; n = 4 mice/group.

To confirm these findings, we next used CSF1R-mer-iCre-mer mice crossed to R26-LSL-tdTomato, which allows for specific labeling of embryonic progenitors and their lineage when tamoxifen is administered during embryonic development. A single dose of tamoxifen was administered in pregnant dams on embryonic day 9.5 (E9.5). Known yolk sac derived cells, brain microglia (Ginhoux *et al*., 2010), showed a high level of recombination near 80% in offspring (**Fig. 2f,g**). However, Ly6C^Hi^ monocytes showed a minimal labeling of tdTom (**Fig. 2g**). Interestingly, nearly 10% of pancreas macrophages displayed tdTomato labeling, demonstrating a subpopulation of pancreas macrophages are derived from embryonic progenitors. Further, quantitation of tdTomato^+^ macrophages showed that after cerulein treatment, this population expanded >3-fold as measured by both flow cytometry and IHC staining (**Fig. 2h-j**).

Importantly, it has been established that embryonically derived macrophages can undergo replacement by monocyte-derived macrophages (MDMs) to varying degrees in a tissue-specific manner (Ginhoux and Guilliams, 2016; Liu *et al*., 2019). Some populations of embryonic macrophages, such as heart macrophages and lung alveolar macrophages undergo gradual replacement (Bain *et al*., 2016; Ginhoux and Guilliams, 2016; Liu *et al*., 2019). Given that the above lineage tracing models are specific for embryonically-derived macrophages, we next sought to measure the total tissue-resident macrophage population in the pancreas irrespective of cellular origin. To accomplish this, we conducted tamoxifen pulse-chase experiments using CSF1R-mer-iCre-mer;LSL-tdTomato adult mice by injecting tamoxifen for 5 consecutive days, then allowing a 10-week chase period for monocytes and short-lived MDMs to turn over and lose tdTomato labeling (**Fig. 2k**). Mice were then treated with vehicle or cerulein, and tdTomato^+^ TRMs were quantitated by flow cytometry and IHC. In agreement with previous embryonic labeling experiments, the number of tdTomato^+^ TRMs was drastically increased with cerulein treatment (**Fig. 2k-n**). Interestingly, while CX3CR1-CreERT2;LSL-tdTomato mice showed similar numerical increases in TRMs when conducting tamoxifen pulse-chase experiments, we maximally observed only 50% of pancreas macrophages labeled by tdTomato (**Supp. Fig. 2e-i**), as others have reported (Calderon *et al*., 2015). To confirm this, we profiled CX3CR1-GFP mice, and while ∼99% of blood monocytes were labeled with GFP, only 45% of pancreas macrophages were GFP^+^ (**Supp. Fig. 2j-k**). This is likely due to differential expression of CX3CR1 gene in pancreas macrophages. IHC staining of CX3CR1-CreERT2;LSL-tdTomato mice further showed that macrophages in the islets of Langerhans were predominantly CX_3_CR1^+^ TRMs (**Supp. Fig. 2l**). As our flow cytometry preparations digest the entire pancreas, we could only distinguish islet macrophages by IHC. We then confirmed islet macrophages are marked by tdTomato in CSF1R-mer-iCre-mer;LSL-tdTomato mice pulse-chased with tamoxifen (**Supp. Fig 2m**), suggesting that islet macrophages are indeed long-lived resident cells marked by CX_3_CR1, as previously described (Calderon *et al*., 2015). Staining these tissues for macrophage marker Lyve1 also showed that islet macrophages lack LYVE1 expression (**Supp. Fig. 2l-m**). Taken together, these data suggest that TRMs in the pancreas consist of both embryonic-derived and adult HSC-derived cells, and both of these TRMs expand numerically during tissue damage. This TRM expansion of cells accompanies an increase in monocyte infiltration and accumulation of MDMs.

### Transcriptional profiling of pancreas macrophages reveals distinct phenotype of tissue-resident and monocyte-derived macrophages

Several studies have shown that ontogeny of TRMs can influence their phenotype and specific function (Bowman et al., 2016; Casanova-Acebes *et al*., 2021; Loyher *et al*., 2018; Zhang *et al*., 2021; Zhu *et al*., 2017). To investigate the ontogeny-dependent roles of macrophages during pancreas injury by pancreatitis or tumor formation, we first performed RNAseq analysis of macrophages sorted from Flt3-YFP mice treated with either vehicle, acute cerulein, or orthotopically implanted PDAC tumors. Differential gene expression testing revealed roughly >2500 differentially expressed genes (DEGs) between Flt3-YFP^+^ and Flt3-YFP^(-)^ macrophages within each treatment condition (**Fig. 3a**). Gene set enrichment analysis (GSEA) showed that genes upregulated in Flt3-YFP^(-)^ macrophages were significantly enriched for pathways related to extracellular matrix (ECM) remodeling and collagen-containing ECM. Conversely, genes upregulated in Flt3-YFP^+^ macrophages were highly enriched for antigen presentation, T cell activation, and lymphocyte co-stimulation pathways (**Fig. 3b**). Flt3-YFP^(-)^ highly expressed genes related to ECM remodeling, growth factors, and scavenger receptors as well as genes commonly used as a resident or alternatively-activated macrophage markers (*Lyve1, Cd163, Siglec1, Cd209, Mrc1*, etc.), while Flt3-YFP^+^ macrophages expressed high levels of antigen-processing and presentation genes such as major histocompatibility complex class II (MHCII) family genes and costimulatory molecules (**Fig. 3c**). Importantly, these differences were retained in both the homeostatic pancreas, tissue injury by pancreatitis and in tumor-bearing pancreas (**Fig. 3c**). These data show, on average, embryonic tissue resident macrophages have district phenotypes with likely enhanced tissue remodeling properties that are retained during tissue injury. Notably, these phenotypes are similar to reports in other tissues (Casanova-Acebes *et al*., 2021; Dick *et al*., 2022; Zhang *et al*., 2021; Zhu *et al*., 2017).

**Figure 3:**
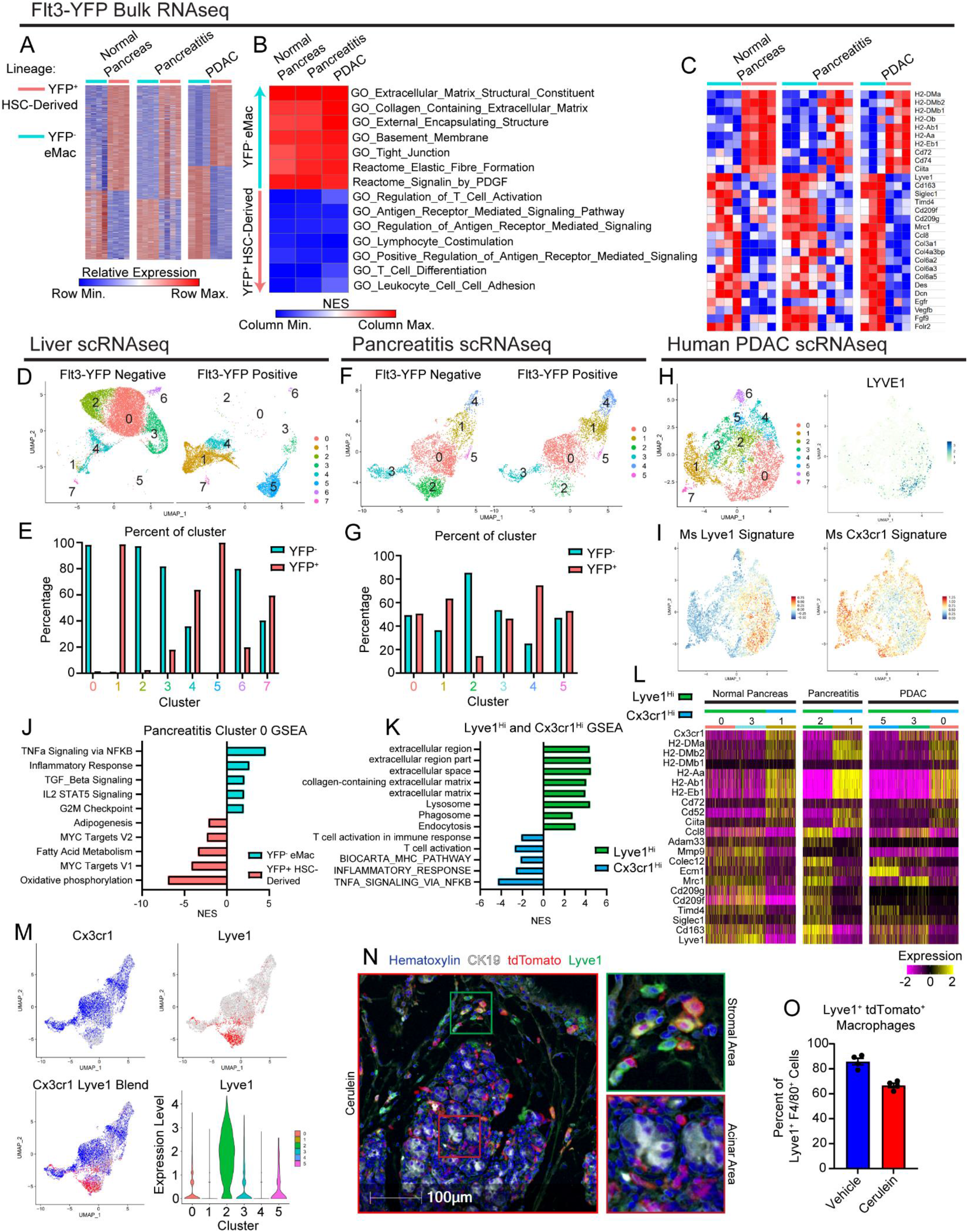
Transcriptional profiling of pancreas macrophages reveals distinct phenotype of tissue-resident and monocyte-derived macrophages. **A**. Heatmap displaying differentially expressed genes in bulk RNAseq data between Flt3-YFP positive and negative macrophages under normal pancreas, pancreatitis, and PDAC conditions. **B**. Heatmap displaying normalized enrichment score of significantly enriched gene sets in Flt3-YFP positive and negative macrophages, pathways selected by FDR < 0.05. **C**. Heatmap displaying differentially expressed genes between Flt3-YFP positive and negative macrophages shared across normal pancreas, pancreatitis, and PDAC samples. **D**. UMAP plot from single-cell RNA-sequencing (scRNAseq) analysis of macrophages sorted from livers hemispleen implanted with KP2 pancreatic cancer cell line. **E**. Quantification of Flt3-YFP lineage by cluster from UMAP in D, displayed as percentage of each cluster. **F**. UMAP plot of scRNAseq analysis of macrophages sorted from cerulein treated pancreas. **G**. Quantification of Flt3-YFP lineage by cluster from UMAP in F, displayed as percentage of each cluster. **H**. UMAP plot of scRNAseq analysis on human monocytes/macrophages from human PDAC dataset, and LYVE1 gene expression. **I**. UMAP plot displaying mouse Lyve1^Hi^ or Cx3cr1^Hi^ macrophage scRNAseq signature using top 100 DEGs from each population mapped into human PDAC data set. **J**. Bar graph of NES values of gene sets comparing Flt3-YFP positive to negative cells within pancreatitis cluster 0 (from F). **K**. Bar graph of NES values of gene sets comparing Lyve1^Hi^ to Cx3cr1^Hi^ macrophages from F. **L**. Heatmap of DEGs upregulated in either Lyve1Hi or Cx3cr1Hi macrophages across normal pancreas, pancreatitis, and PDAC samples. **M**. UMAP plots displaying Cx3cr1, Lyve1, or merged gene expression from pancreatitis macrophages (from F) and violin plot showing Lyve1 gene expression by cluster of pancreatitis macrophages. **N**. Representative images from multiplex immunohistochemistry (mIHC) staining of pancreas tissue from vehicle or cerulein treated CSF1R-mer-iCre-mer;LSL-tdTomato tamoxifen pulse/chase mice. Tissue stained for Hematoxylin, F4/80, tdTomato, Lyve1, and CK19. **O**. Quantification of percentage of Lyve1^+^ macrophages labeled by tdTomato (Lyve1^+^ tdTomato^+^ macrophages as percentage of total Lyve1^+^ macrophages); n = 4 mice/group. Data are presented as mean ± SEM unless otherwise indicated. n.s., not significant; *p <0.05. For comparisons between two groups, Student’s two-tailed t-test was used, except for (A),(C),(L), where Bonferroni correction was used, and (B),(J),(K), where FDR was used.

We next wanted to determine if embryonically-derived TRMs defined phenotypically unique subpopulations of macrophages during pancreas injury, or if HSC-derived tissue-resident macrophages and embryonic macrophages blended at the level of phenotypic subpopulations. To accomplish this, we performed single-cell RNA-sequencing (scRNAseq) on sorted Flt3-YFP^+^ and Flt3-YFP^(-)^ macrophages from homeostatic pancreas, pancreatitis, and PDAC tumors in the pancreas. Additionally, we analyzed tumor bearing liver and lung as controls, because liver Kupffer cells would serve as a positive control for a distinct embryonic tissue resident macrophage population with little contribution from monocytes while lung alveolar macrophages would be gradually replaced with monocytes (Ginhoux and Guilliams, 2016; Liu *et al*., 2019; Schulz *et al*., 2012; Yona *et al*., 2013). As expected, even under tumor conditions, liver and lung macrophage populations were from distinct origins. In a model of PDAC metastasis to the liver, *Clec4f* expressing Kupffer cells (cluster 0 & 2, **Fig. 3d, Supp. Fig. 3h**) were almost exclusively derived from Flt3-YFP^(-)^ cells, consistent with Kupffer cells being strictly embryonically derived (**Fig. 3d-e**). Similarly, alveolar macrophages from lungs bearing KP tumors were >60% derived from Flt3-YFP^(-)^ embryonic origins (cluster 0, **Supp. Fig. 3i-j**). Conversely, lung interstitial macrophages were exclusively observed as being Flt3-YFP^+^ (cluster 1 & 3, **Supp. Fig. 3i-j**). This contrasts with what we observed in homeostatic pancreas, pancreatitis, and PDAC tumors, where macrophage populations were of much more mixed embryonic and HSC-derived origin (**Fig. 3f-g**). Across homeostatic and pathologic states, most macrophage clusters had contribution from both Flt3-YFP^+^ and YFP^(-)^ macrophages, suggesting clusters were not completely defined by developmental origin. The notable exceptions to this were clusters highly expressing *Lyve1* (homeostasis cluster 0, pancreatitis cluster 2, and PDAC cluster 5, **Supp. Fig. 3b-g**), which across treatment conditions showed enrichment for Flt3-YFP^(-)^ macrophages of embryonic origin.

The highest-level being cluster 2 in pancreatitis samples that were 90% YFP^(-)^ embryonically derived (**Fig. 3f-g**). Conversely, some clusters showed bias towards Flt3-YFP^+^ sources and were marked with *Cx3cr1* expression and high levels of MHCII family genes (homeostasis clusters 2 & 4, and pancreatitis cluster 4). Selective expression of *Cx3cr1* gene across macrophage clusters further supports previous lineage tracing data using CX3CR1-CreERT2 mice (**Supp. Fig. 2d-i**). Some Cx3cr1^+^ clusters also likely represent TRMs of mixed origin (Chakarov *et al*., 2019; Liu *et al*., 2019). By example cluster 0 in pancreatitis samples was nearly 50% comprised of Flt3-YFP^+^ cells (**Fig. 3f-g**) and was marked by *Cx3cr1*, some MHCII family genes, *Apoe*, and *Trem2*, but notably lacked *Ccr2* expression, which has been used in other tissues such as the heart to delineate some TRM and MDM populations (Epelman *et al*., 2014a; Epelman et al., 2014b). These clusters highlight the ability of monocyte-derived cells to gradually replace embryonically derived macrophages and become long-lived TRMs, but will still be marked as Flt3-YFP^+^. Taken together, this suggests that aside from Lyve1^Hi^ TRMs, most macrophage populations come from mixed origin and unlike observations in liver and lung tissue, macrophage origin in the pancreas does not delineate every cluster of phenotypic macrophages, which may be defined more by time in residence and subregional spatial contexture within the tissue.

To investigate if developmental origin still contributed phenotypic biases within subpopulations of potential TRM clusters, we compared Flt3-YFP^+^ cells to Flt3-YFP^(-)^ cells within cluster 0. Unlike previous observations in bulk RNAseq data, this comparison did not show enrichment in ECM remodeling gene sets in Flt3-YFP^(-)^ cells, nor enrichment in antigen presentation gene sets in Flt3-YFP^+^ cells. Instead, Flt3-YFP^(-)^ cells were enriched for TNFα signaling by NF-κB, inflammatory responses, and TGF-β signaling, while Flt3-YFP^+^ cells showed higher expression of oxidative phosphorylation and Myc targets (**Fig. 3j**). Additionally, as a positive control to determine if similar ECM remodeling or antigen presentation gene sets were detected in our scRNAseq data, we compared all Flt3-YFP^+^ to Flt3-YFP^(-)^ macrophages irrespective of cluster. Indeed, this showed similar enrichment of ECM remodeling and collagen gene sets in Flt3-YFP^(-)^ macrophages, while Flt3-YFP^+^ had enrichment in T cell proliferation and immunity, and antigen processing and presentation (**Supp. Fig. 3a**). Taken together, these data suggest developmental origin does yield phenotypic biases on both the bulk macrophage level and inside of phenotypic subsets; but within a macrophage subset tissue residence and not developmental origin is dominant for tissue remodeling vs antigen presentation phenotypes. This agrees with prior studies in other tissues during homeostasis (Lavin *et al*., 2014; Wang et al., 2020).

Investigating gene expression phenotypes of macrophage clusters, we noted macrophages fell into LYVE1^Hi^MHCII^Lo^ CX_3_CR1^Lo^ and LYVE1^Lo^ MHCII^Hi^ CX_3_CR1^Hi^ subpopulations. These phenotypes fit with other reports for TRMs in fat, dermis, heart, lung, mesenteric membranes, and other tissues (Chakarov *et al*., 2019; Dick *et al*., 2022; Zhang *et al*., 2021). Consistent with LYVE1^Hi^ macrophages being predominantly Flt3-YFP^(-)^, these clusters had notably similar gene expression profiles as was observed in bulk RNAseq of Flt3-YFP^(-)^ macrophages. These clusters had expression of scavenger receptors and alternative-activation markers such as *Cd163, Siglec1, Timd4, Cd209*, and *Mrc1*. LYVE1^Hi^ clusters also showed enrichment in ECM and collagen-related gene sets, as well as some enrichment in endocytosis and phagocytosis (**Fig 3k-l**). Similarly, CX_3_CR1^Hi^ clusters expressed the same phenotype as was observed in bulk RNAseq of Flt3-YFP^+^ macrophages, with high expression of antigen-presentation and costimulatory genes, including *H2-Aa, H2-Ab1, H2-Eb1, Cd52*, and *Cd74*. GSEA analysis of CX_3_CR1^Hi^ macrophages showed enrichment in T cell activation, MHC pathways, as well as inflammatory and NF-κB signaling. This further suggests that while clusters or populations of macrophages have varying degrees of enrichment in their cellular origin, there are correlations in phenotypes; with LYVE1^Hi^ being mostly Flt3-YFP^-^ and involved in remodeling ECM, while CX_3_CR1^Hi^ are more Flt3-YFP^+^ and involved in antigen processing and presentation.

We next examined how closely these populations correlated with macrophages in human pancreas and PDAC scRNAseq datasets (Zhou et al., 2021). Human PDAC macrophage and monocyte clusters were identified, isolated, and re-clustered, revealing eight total clusters (**Fig 3h**). Gene expression signatures of LYVE1^Hi^MHCII^Lo^ and LYVE1^Lo^MHCII^Hi^ macrophages from our mouse dataset were mapped into human scRNAseq data using an averaged z-score (**Fig 3h-i**). The LYVE1^Hi^MHCII^Lo^ signature was most highly expressed in cluster 0, which was noted to be the only cluster with *LYVE1* gene expression (**Fig 3i**). Similarly, the LYVE1^Lo^MHCII^Hi^ signature highlighted a few clusters, but was highest in cluster 1, which also expressed *CCR2* and is likely a monocyte-derived macrophage population (**Fig 3h-i**). These data suggest the identified human macrophage subsets share similar gene expression phenotypes as mouse LYVE1^Hi^MHCII^Lo^ and CX_3_CR1^Hi^MHCII^Hi^ macrophages.

As the Flt3-YFP lineage tracing system specifies developmental origin, but does not distinguish time in tissues, we next wanted to determine if LYVE1^Hi^ macrophages were predominantly made up of long-lived resident cells. To answer this question, we accessed tissue from pulse-chase experiments in CSF1R-mer-iCre-mer;LSL-tdTomato mice (**Fig 2**). We then performed multiplex IHC (mIHC) staining on pancreas tissue of these mice for F4/80, LYVE1, tdTomato, and CK19 to quantitate the percentage of LYVE1^Hi^ macrophages that were tissue-resident (tdTomato^+^). A majority (>80%) of F4/80^+^LYVE1^Hi^ cells were labeled by tdTomato and identified as resident cells in homeostatic pancreas. Analysis of the localization of these LYVE1^+^ TRMs showed these cells were located between lobules of the pancreas in stromal areas and on the very surface of the pancreas tissue and rarely in close contact with acinar cell clusters or within islets (**Fig 3n**). This distribution was not mirrored by all tdTomato+ TRMs, some of which showed infiltration between clusters of acinar cells suggesting that LYVE1^Hi^MHCII^Lo^ and LYVE1^Lo^MHCII^Hi^ TRMs have distinct anatomical distribution within the pancreas depending on their phenotypic role. To assess this unique distribution of LYVE1^+^ macrophages in humans, we performed mIHC for CD163, LYVE1, and CK19 on human PDAC, pancreatitis, or adjacent normal pancreas tissues. Similar to mouse models, in human pancreatitis and PDAC tissues, we only observed LYVE1^+^CD163^+^ macrophages within the stroma and outside of acinar areas of the pancreas, while LYVE1^(-)^CD163^+^ macrophages were localized in both compartments (**Supp. Fig. 3l-n**). While the desmoplastic stromal area is limited in adjacent normal pancreas samples, LYVE1^+^CD163^+^ macrophages were located between lobules of the pancreas and on the border of the tissue, similar to mouse tissues. Taken together, these data suggest that LYVE1^+^ TRMs have both the phenotype and localization to play a role in regulating pathologic fibrosis during pancreas injury and tumorigenesis.

### Tissue-resident macrophages are required for maintaining tissue integrity during pancreatitis

We next asked whether TRMs or MDMs have a distinct functional impact on pancreas injury by pancreatitis. To deplete TRMs, we used a combination of colony-stimulating factor-1 (CSF1) neutralizing antibodies and clodronate-loaded liposomes (denoted as αCSF1-CLD herein), as previously reported, (Yu et al., 2021; Zhu *et al*., 2017). Mice were treated with a combination of αCSF1 and CLD, then given a 10-day recovery period to allow for recovery of blood monocyte and MDM numbers. Following the 10-day recovery period, blood monocyte and pancreas macrophage numbers were quantified by flow cytometry. As expected, there was no difference in Ly6C^Hi^ monocyte numbers between αCSF1-CLD and IgG-PBS treated mice after the recovery period, but notably, Ly6C^Lo^ monocytes are slower to recover (**Fig. 4b-c**). Immediately following the recovery period (Day -1), total pancreas macrophage number was significantly reduced (**Fig. 4d**), and MHCII^Lo^ macrophage subset, noted above to be enriched for TRMs, was the most significantly depleted subset (**Supp. Fig. 4c**). Following recovery period, mice were then treated with cerulein for 7 days, and again pancreas macrophage number was quantified. IgG-PBS treated mice showed similar accumulation of macrophages as seen before, however αCSF1-CLD treatment, in spite of having no detriment in Ly6C^Hi^ monocytes, was able to effectively restrain accumulation and total number of macrophages in the injured pancreas (**Fig. 4d**). Similarly, this depletion was most dramatic in the MHCII^Lo^ subset of macrophages (**Supp. Fig. 4b-c**). Taken together, these data demonstrate that not only is αCSF1-CLD treatment an effective TRM-depletion strategy, but also highlighting the more substantial increase in TRMs compared to MDMs.

**Figure 4:**
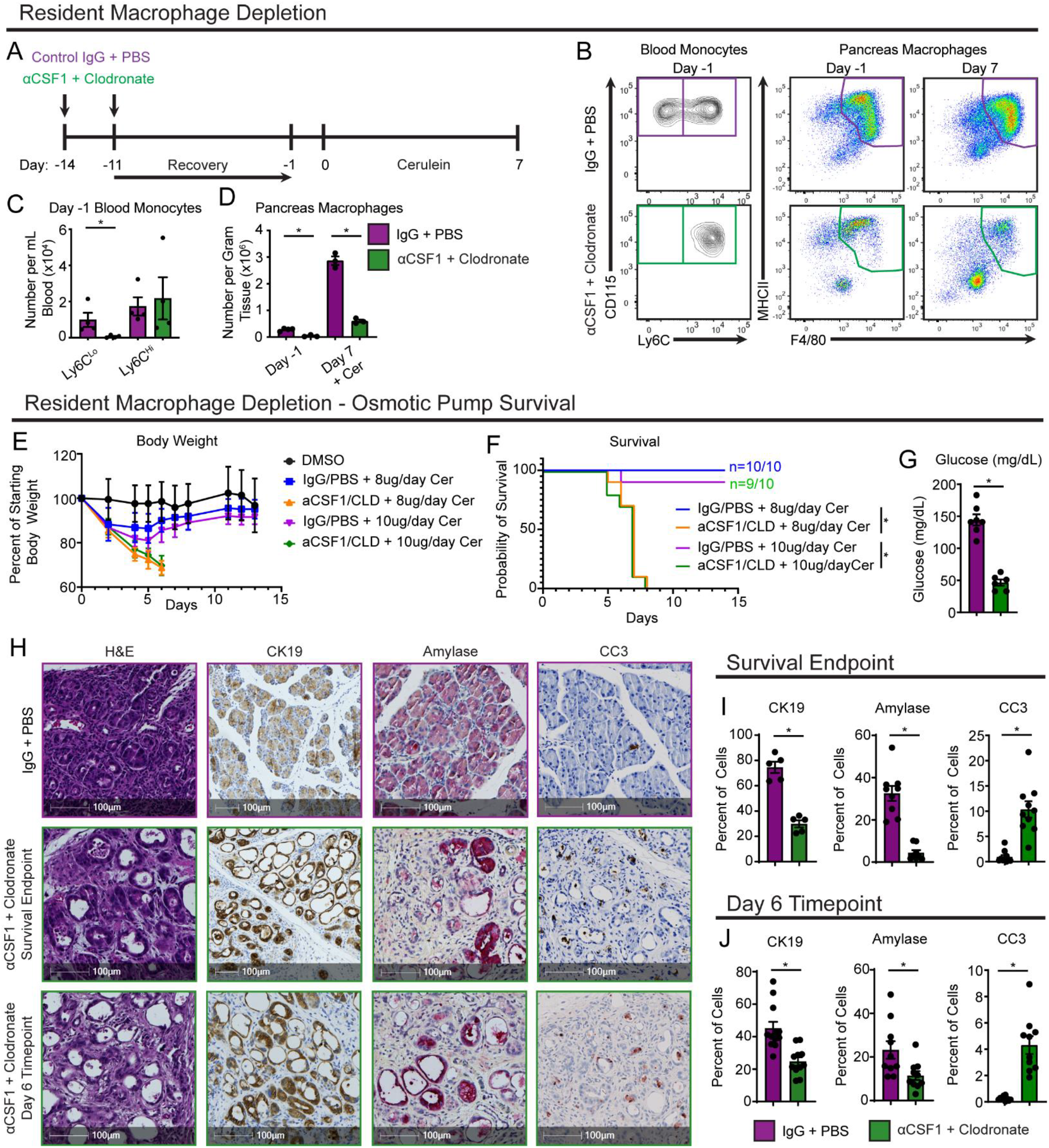
Tissue-resident macrophages are required for maintaining tissue integrity during pancreatitis. **A**. Treatment scheme for resident macrophage depletion with αCSF1-CLD. **B**. Flow cytometry plots of blood monocytes and pancreas macrophages after recovery period (day -1) and after cerulein treatment (day 7) with and without αCSF1-CLD depletion. **C**. Quantification of blood Ly6C^Lo^ and Ly6C^Hi^ monocytes after recovery period (day -1). **D**. Quantification of pancreas macrophages after recovery period (day -1) and after cerulein treatment (day 7) with and without αCSF1-CLD depletion. **E**. Mouse body weight measurement following implantation of osmotic pump for cerulein delivery, with and without αCSF1-CLD depletion. 8ug/day and 10ug/day cerulein concentration as indicated. **F**. Kaplan-Meier survival curve showing IgG-PBS and αCSF1-CLD groups. 8ug/day and 10ug/day cerulein concentration as indicated. **G**. Blood glucose concentration taken at survival endpoint in IgG-PBS and αCSF1-CLD mice with 10ug/day cerulein treatment. **H**. Representative images of pancreas tissue stained for H&E, CK19, Amylase, and CC3 from mice treated with IgG-PBS or αCSF1-CLD and implanted with cerulein-loaded osmotic pumps until survival endpoint (top two rows) or for 6 days (bottom row). **I**. Quantification of CK19, Amylase, and CC3 IHC stains at cerulein treatment survival endpoint displayed as the percentage of cells; n = 5-10 mice/group. **J**. Quantification of CK19, Amylase, and CC3 IHC stains at day 6 cerulein treatment timepoint displayed as the percentage of cells; n = 10-12 mice/group. Data are presented as mean ± SEM unless otherwise indicated. n.s., not significant; *p <0.05. For comparisons between two groups, Student’s two-tailed t-test was used.

To investigate the potential functional outcomes that TRMs may have on tissue damage, mice were implanted with osmotic pumps to continually administer cerulein following TRM depletion for up to 14 days. In control mice, we observed rapid acute body weight loss, followed by recovery, and after 14 days mice were gaining body weight and analysis pancreas tissue showed almost complete recovery (**Fig. 4e-h**). By contrast, mice depleted of TRMs with CSF1-CLD had progressive bodyweight loss, and 100% of mice either died or reached our humane survival endpoints (>20% weight loss) by 8 days post pancreatitis induction (**Fig. 4e-f**). These observations were consistent across two cerulein doses and in both C57BL/6 and FvBn genetic backgrounds (**Fig. 4e-f, Supp. Fig. 4g-i**). Consistent with loss of pancreas function, TRM-depleted mice also had significant decreases in blood glucose levels (**Fig. 4g**) and higher serum amylase (**Supp. Fig. 4e**), both of which are hallmarks of severe pancreatitis. Histological analysis of pancreas tissue at survival endpoint also confirmed loss of pancreas tissue integrity, marked ADM, and significant tissue necrosis in TRM-depleted mice, compared to non-depleted controls which looked relatively healthy (**Fig. 4h**). IHC analysis at survival endpoint showed TRM-depleted mice had a greater than 90% loss of amylase^+^ acinar cells, as well as a significant loss of CK19^+^ ductal cells, and both cell types had significant levels of apoptosis marked by cleaved-caspase-3 (CC3^+^) (**Fig. 4i**). Similar changes were observed at a six-day post cerulein time point analysis, comparing TRM-depleted mice to controls (**Fig. 4j**). Finally, to rule out TRM-depletion causing potential liver damage, liver tissues were stained by H&E, and no histologic difference was noted between αCSF1-CLD and IgG-PBS groups (**Supp. Fig. 4f**). We next wanted to conversely deplete MDMs, to determine if they confer any protective role. To impair MDMs, we utilized CCR2-knockout (CCR2-KO) mice, as CCR2 deficiency impairs the egression of Ly6C^Hi^ monocytes from the bone marrow (Serbina and Pamer, 2006). In contrast to TRM-depletion, CCR2-KO mice were highly reduced in Ly6C^Hi^ blood monocytes (**Supp. Fig. 4i**), however this did not translate to a change in either total macrophage number or MHCII^Hi^ macrophage number under steady-state (vehicle-treated) conditions (**Supp. Fig. 4j-m**). Following cerulein treatment, total macrophage number was again not significantly altered in CCR2-KO mice (**Supp. Fig. 4k**), however, there was a decrease in the accumulation of MHCII^Hi^ macrophages (**Supp. Fig. 4l**). The fact that the decrease in MHCII^Hi^ MDMs does not largely affect the total number of macrophages that accumulate in the pancreas again highlights that TRMs expand more significantly than MDMs with cerulein treatment. Interestingly, when implanted with osmotic pumps to deliver cerulein, CCR2-KO mice showed no difference in survival, change in body weight, or pancreas weight compared to CCR2-wildtype controls (**Supp. Fig. 4n-o**). Taken together, these data suggest that TRMs are critically important in restoring tissue homeostasis in the pancreas following tissue damage by pancreatitis, whereas MDMs are largely dispensable.

### Depletion of tissue-resident macrophages does not alter ADM induction, but rather attenuates the fibrotic response

We next sought to understand how TRMs might be influencing the pancreas microenvironment and restraining tissue damage. We first asked whether the protective effects that TRMs have on restraining tissue damage were caused by somehow altering the ADM process. The ability for acinar cells to go through the ADM process and re-enter the cell cycle is critical for regenerating cells to heal the pancreas following injury (Jensen et al., 2005; Mills and Sansom, 2015; Willet et al., 2018). For these studies, we returned to serial IP injection of cerulein (**Fig. 1c-d**), which allows for improved mouse survival even after TMR-depletion. Interestingly, αCSF1-CLD treated mice were observed to have increased percentage of total cells expressing CK19 when analyzed as a single IHC stain (**Fig. 5a**). However, closer examination of the pancreas histology revealed that TRM-depleted mice had significantly fewer stromal cells and stromal area accumulating between acinar cell clusters, which perhaps convoluted this CK19 analysis (**Fig 5a**). To better profile potential changes in ADM, we developed a multiplex-IHC (mIHC) panel that would allow us to quantify ADM markers CK19, Sox9, and Ki-67 within amylase-expressing acinar cells. As opposed to previous single-plex CK19 analyses, mIHC analysis showed no difference in CK19, Sox9, or Ki-67 as a percentage of acinar cells (**Fig. 5b-c**). This suggests that TRM-depletion does not affect the ability for acinar cells to undergo the ADM process, but rather alters survival after ADM, as was observed previously with TRM-depletion causing a significant increase in apoptotic acinar cells (**Fig 4h-j**). We next hypothesized that acinar cell survival could be linked to protective effects of stromal cells that accumulate during pancreatitis and assessed changes in fibroblasts by podoplanin (PDPN) staining. Indeed, TRM-depleted mice showed a >50% reduction in fibroblast numbers (**Fig. 5b-c**). Taken together, these data suggest that while TRMs do not dramatically regulate acinar cell progression through the ADM process, they do affect their survival by promoting protective stromagenesis.

**Figure 5:**
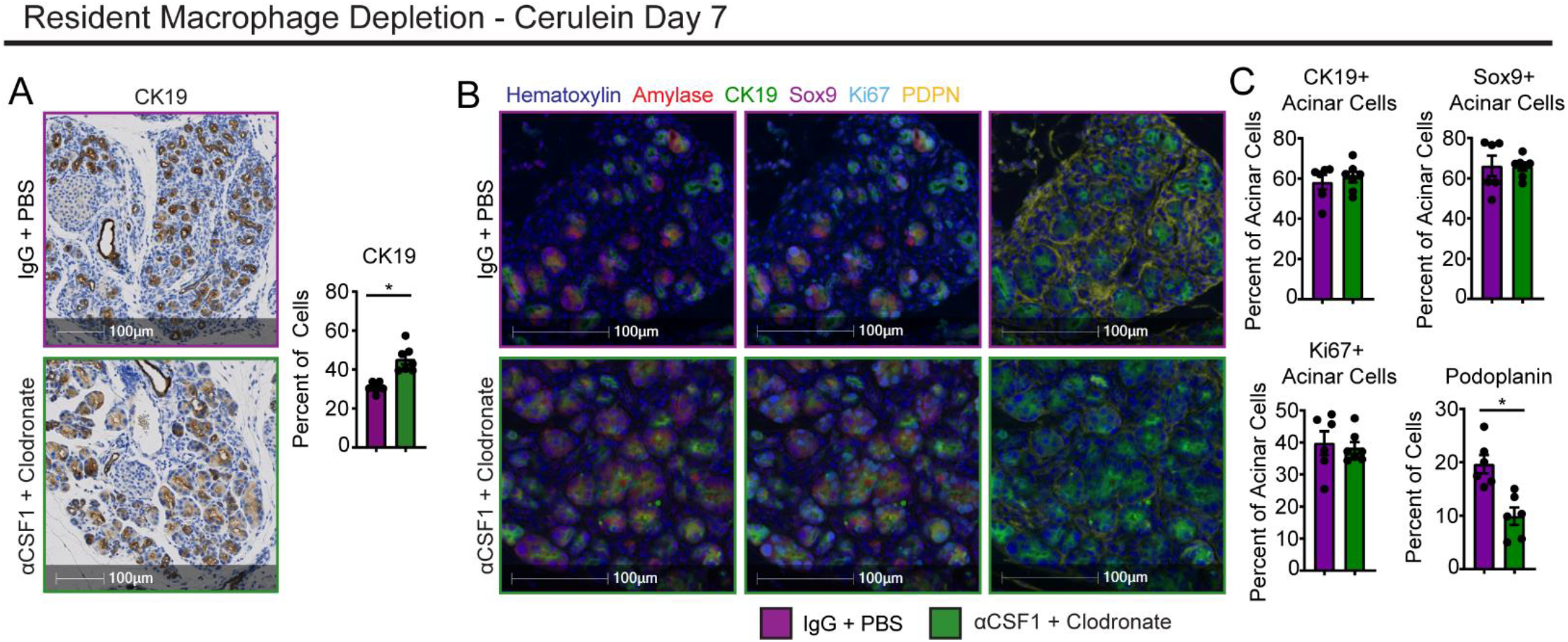
Depletion of tissue-resident macrophages does not alter ADM induction, but rather attenuates the fibrotic response. **A**. Representative images and quantification of CK19 IHC stain on IgG-PBS and αCSF1-CLD pancreas tissue treated with cerulein by serial I.P. injection; n = 5-7 mice/group. **B**. Representative images of mIHC staining of IgG-PBS and αCSF1-CLD pancreas tissue treated with cerulein by serial I.P. injection. Stain includes Hematoxylin, Amylase, CK19, Sox9, Ki-67, and podoplanin (PDPN). **C**. Quantification of percentage of acinar cells positive for CK19, Sox9, or Ki-67, and percentage or total cells positive for podoplanin; n = 6-7 mice/group. Data are presented as mean ± SEM unless otherwise indicated. n.s., not significant; *p <0.05. For comparisons between two groups, Student’s two-tailed t-test was used.

### Tissue-resident macrophages are required for shaping the tissue-protective fibrotic response

To further profile how TRMs are involved in driving the fibrotic response during pancreatitis, we examined fibroblast and ECM changes across pancreatitis progression. For these studies, we depleted TRMs with αCSF1-CLD treatment as before, then induced pancreatitis for varying lengths of time to simulate early initiation of disease, acute disease phase, or chronic disease. Across all time points, TRM-depleted mice had reduced fibroblasts measured by PDPN staining (**Fig. 6a-b, Supp. Fig. 6a-b**), confirming that TRMs are likely causing accumulation of fibroblasts. In early time points (3 and 7 days cerulein), the critical ECM molecule fibronectin was only observed to be mildly reduced. However, by day 17 of cerulein treatment, there was a considerable reduction in fibronectin deposition (**Fig. 6a-b, Supp. Fig. 6a-b**). As an additional control to determine if αCSF1-CLD treatment could directly deplete fibroblasts, steady-state pancreas tissue was taken from αCSF1-CLD and IgG-PBS treated mice, and showed no difference in PDPN staining (**Supp. Fig. 6c**). Next, to confirm these changes and profile fibroblast subpopulations, we quantified total fibroblasts, as well as four distinct subsets that have been previously described in for homeostatic pancreas and PDAC (Elyada et al., 2019; Öhlund et al., 2017); Ly6C^+^ inflammatory fibroblasts (iFibs), MHCII^+^ antigen-presenting fibroblasts (apFibs), alpha-smooth muscle actin (αSMA) expressing myofibroblasts (myFibs), and a triple-negative (Ly6C^(-)^MHCII^(-)^αSMA^(-)^) subset marked only by platelet-derived growth factor receptor alpha (PDGFRα), referred to as PDGFRα^+^ fibroblasts herein (**Supp. Fig. 6d**). Nomenclature for fibroblast subsets was also adapted from previous studies of PDAC (Elyada *et al*., 2019; Öhlund *et al*., 2017). In flow cytometric analysis, total fibroblasts were markedly increased during pancreatitis in control IgG-PBS mice (**Fig. 6c-d**). However, fibroblast numbers failed to expand in αCSF1-CLD treated mice (**Fig. 6c-d**). Analysis of fibroblast subsets observed a similar increase in Ly6C^+^ and PDGFRα^+^ fibroblasts during pancreatitis, and failure of these subsets to expand in TRM-depleted mice (**Fig. 6d**). Interestingly, unlike PDAC, we observed very few αSMA^+^ and MHCII^+^ fibroblasts, and they did not expand dramatically by 7 days of pancreatitis, suggesting these populations are defined either by prolonged tissue damage or tumor-specific conditions (**Supp. Fig. 6e**).

**Figure 6:**
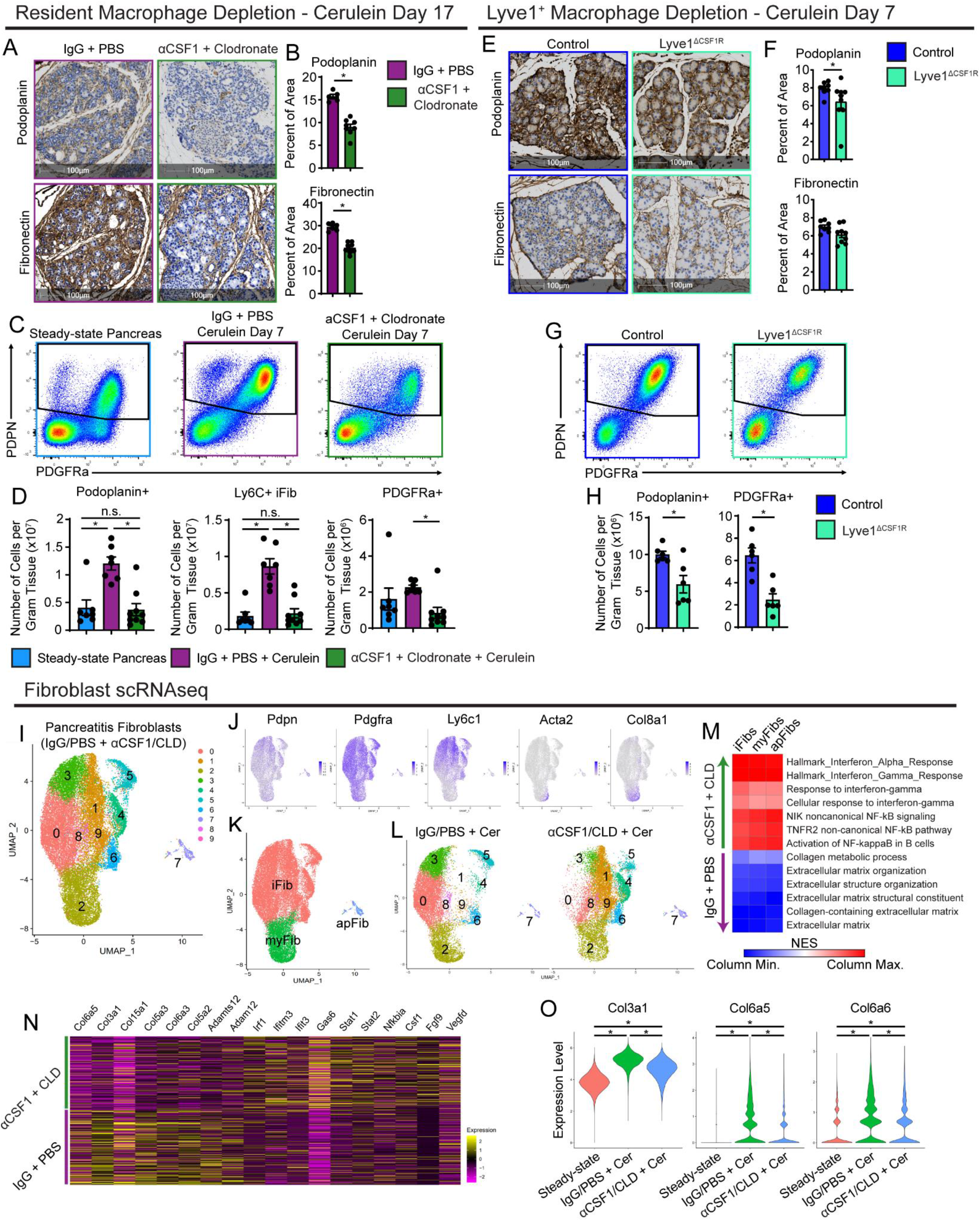
Tissue-resident macrophages are required for shaping the tissue-protective fibrotic response. **A**. Representative images of pancreas tissue stained for podoplanin or fibronectin, treated with either IgG-PBS or αCSF1-CLD and cerulein by serial I.P. injection for 17 days. **B**. Quantification of podoplanin and fibronectin area; n = 6-8 mice/group. **C**. Flow cytometry plots of fibroblasts under steady-state pancreas, IgG-PBS and cerulein, or αCSF1-CLD and cerulein conditions. **D**. Density of total podoplanin^+^ fibroblasts, Ly6C^+^ iFibs, and PDGFRa^+^ fibroblasts under steady-state pancreas, IgG-PBS and cerulein, or αCSF1-CLD and cerulein conditions; n = 6-9 mice/group. **E**. Representative images of pancreas tissue stained for podoplanin or fibronectin from control (Lyve1-Cre^(-)^ mice) or Lyve1-Cre;CSF1R^flox/flox^ mice (Lyve1^ΔCSF1R^) treated with cerulein. **F**. Quantification of podoplanin and fibronectin positive areas; n = 8-9 mice/group. **G**. Flow cytometry plots of fibroblasts from control and Lyve1^ΔCSF1R^ mice. **H**. Density of total podoplanin+ fibroblasts and PDGFRa+ fibroblasts from control and Lyve1^ΔCSF1R^ mice; n = 6 mice/group. **I**. UMAP plot of pancreas fibroblasts sorted from IgG-PBS and αCSF1-CLD mice treated with cerulein. **J**. Gene expression of fibroblast marker genes Pdpn, Pdgfra, Ly6c1, Acta2, and Col8a1. **K**. UMAP plot showing the classification of fibroblast clusters into iFib, myFib, and apFib subtypes. **L**. UMAP plot split by sample (IgG-PBS and αCSF1-CLD). **M**. Heatmap displaying NES of gene sets significantly enriched across fibroblast subtypes in IgG-PBS or αCSF1-CLD samples. **N**. Heatmap displaying gene expression of select differentially expressed genes between IgG-PBS and αCSF1-CLD samples. **O**. Violin plots displaying gene expression of significantly differentially expressed genes across steady-state pancreas, IgG-PBS and cerulein, or αCSF1-CLD and cerulein conditions. Data are presented as mean ± SEM unless otherwise indicated. n.s., not significant; *p <0.05. For comparisons between two groups, Student’s two-tailed t-test was used, except for (O), where Bonferroni correction was used, and (M), where FDR was used.

Given that LYVE1^+^ macrophages represented a major embryonically-derived TRM population (**Fig. 3f-g**,**m**) and were uniquely located within stromal areas of the pancreas (**Fig. 3n**), we next sought to determine if they play a role in initiating the fibrotic response. To answer this question, we crossed Lyve1-Cre mice to CSF1R^flox/flox^ mice (termed Lyve1^ΔCSF1R^), to specifically delete the CSF1R gene in LYVE1-expressing cells. As CSF1R signaling is known to be required for macrophage differentiation and survival, this mouse results in selective Lyve1^+^ macrophage loss, as previously shown for macrophages (Zhang *et al*., 2021). In pancreas tissues, we also observed a significant reduction in LYVE1^+^ macrophages in Lyve1^ΔCSF1R^ mice compared to Lyve1-Cre^(-)^ littermate controls (**Supp. Fig. 6f-g**). Next, we induced pancreatitis in Lyve1^ΔCSF1R^ and controls and found that Lyve1^ΔCSF1R^ mice had reduced fibroblast expansion by both PDPN IHC staining (**Fig. 6e-f**), and flow cytometry analyses for PDPN^+^ fibroblasts and PDGFRα^+^ fibroblasts (**Fig. 6g-h**). Conversely, to check the effect of MDMs on fibroblast expansion, we compared pancreatitis tissues in CCR2-KO mice and controls. Unlike TRM-depletion, we found no differences in pancreas fibroblast numbers or ECM deposition by IHC or flow cytometry during pancreatitis in CCR2-KO mice compared to controls (**Supp. Fig. 6i-l**). Taken together, these data suggest that Lyve1^+^ TRMs coordinate the fibrotic response to retain pancreas tissue integrity and function.

We next sought to understand how TRMs impact fibroblast expansion and phenotype. To accomplish this, we performed scRNAseq after sorting total fibroblasts from αCSF1-CLD and IgG-PBS mice treated with cerulein. Clustering of αCSF1-CLD and IgG-PBS samples resulted in 10 distinct clusters that broadly fit into the iFib, myFib, and apFib subsets based on gene expression (**Fig. 6i-k**). The iFib subset again seemed to be the dominant population, with clusters 0, 1, 3, 4, 5, 6, 8, and 9 having high expression of iFib markers *Ly6c1* and *Clec3b* (**Fig. 6j-k**). Although cluster 2 had limited expression of *Acta2* (previously used to define the myofibroblast subset), they highly expressed *Col8a1* and *Cxcl14*, which others have used to define myofibroblasts (Elyada *et al*., 2019; Öhlund *et al*., 2017). Lastly, cluster 7, although very limited in number, had high expression of antigen-presentation genes *H2-Ab1, Cd74*, and *Slpi*, consistent with an apFib-like identity (Elyada *et al*., 2019; Hosein et al., 2019). Comparing αCSF1-CLD and IgG-PBS samples, there was a clear shift in fibroblast populations (**Fig. 6l**). Clusters 0 and 3 of the iFib subset were almost exclusively represented in the IgG-PBS samples, while iFibs from the αCSF1-CLD samples clustered separately in clusters 1 and 4 (**Fig. 6l**). Because we observed such clear shifts in populations between groups, we next ran differential gene expression testing by comparing all iFibs in the TRM-depleted to iFibs in the IgG-PBS samples. GSEA analysis revealed IgG-PBS samples were enriched in gene sets related to ECM production and organization, and collagen production. Conversely, TRM-depleted fibroblasts were overall more inflammatory, showing enrichment in interferon alpha and gamma responses, as well as NF-κB signaling pathway related gene sets (**Fig. 6m**). These pathways are known drivers of pancreatitis severity (Folias et al., 2014; Habtezion, 2015; Liou *et al*., 2013; Manohar et al., 2017). Enrichment in these gene sets was consistent across all fibroblast subsets, with myFibs and apFibs showing similar changes (**Fig. 6m**). At the gene level across all fibroblasts, IgG-PBS cells indeed showed upregulation of many collagen genes and other ECM remodeling proteins, while fibroblasts from TRM-depleted pancreas showed upregulation in interferon response genes *Irf1, Ifitm3, Ifit3, Gas6*, as well as *Nfkbia*, involved in NF-κB signaling (**Fig. 6n**). Interestingly, when including fibroblasts sorted from homeostatic pancreas in our analysis, these cells clustered separately from, and expressed collagen and ECM genes at much lower levels than, both cerulein treated samples. However, IgG-PBS cells upregulated these genes to a higher level than αCSF1-CLD cells did (**Fig. 6o, Supp. Fig. 6m-o**). These changes suggest that while TRMs are present, fibroblasts highly upregulate genes related to collagen and ECM to contribute to fibrosis, but when TRMs are depleted, fibroblast activation is altered. Fibroblasts are not only reduced in number, but present a more inflammatory phenotype, no longer producing protective fibrosis, but instead promoting inflammation that could be contributing to acinar cell death and worse overall outcomes.

### Tissue-resident macrophages drive fibrosis and pancreatitis-accelerated PDAC progression

Tumor-associated macrophages (TAMs) have previously been implicated in playing tumor-supportive roles in PDAC, and our previous study has shown a large portion of TAMs are embryonically derived and express a similar pro-fibrotic phenotype (Zhu *et al*., 2017). We next wanted to understand if TRMs in the pancreas support tumor growth and survival through a similar mechanism of fibroblast activation and initiating fibrosis. If TRM-induced fibrosis during pancreatitis played a protective role critical for acinar cell survival, we hypothesized that this would similarly facilitate transformed ductal cells growth and survival. To test this hypothesis, we utilized several genetic models of PDAC. First, to study early tumorigenesis, we used an inducible KRAS model (iKRAS*) that expresses KRAS^G12D^ within the pancreas upon administration of doxycycline (doxy, **Supp. Fig. 7a**). Previous studies have shown that brief cerulein administration accelerates transformation and tumor growth (Collins et al., 2012; Guerra et al., 2007). As such, we treated iKRAS* mice with cerulein after TRM-depletion or control treatment, then placed mice on doxy for three weeks (**Supp. Fig. 7a**), after which pancreas histology showed widespread growth of early-stage tumor lesions. TRM-depleted mice showed a significant reduction in pre-malignant pancreas weight (**Fig. 7b**), suggesting that TRMs are involved in supporting neoplastic growth. To understand if TRMs are acting by driving a fibrotic stromal response, pancreas tissue was analyzed by machine-learning software trained to distinguish tumor lesions from stromal areas. This analysis showed slight reduction in the percent of pancreas classified as stroma in αCSF1-CLD treated mice (**Fig. 7c**). As another metric, we stained tissue for CK19 and found it increased as a percentage of the pancreas, consistent with diminishment of induction of desmoplasia in the αCSF1-CLD treated group (**Fig. 7d**). Taken together, these data support that TRMs similarly facilitate the expansion of the fibrotic stroma during tumor development, and that when depleted, tumors grow more slowly.

**Figure 7:**
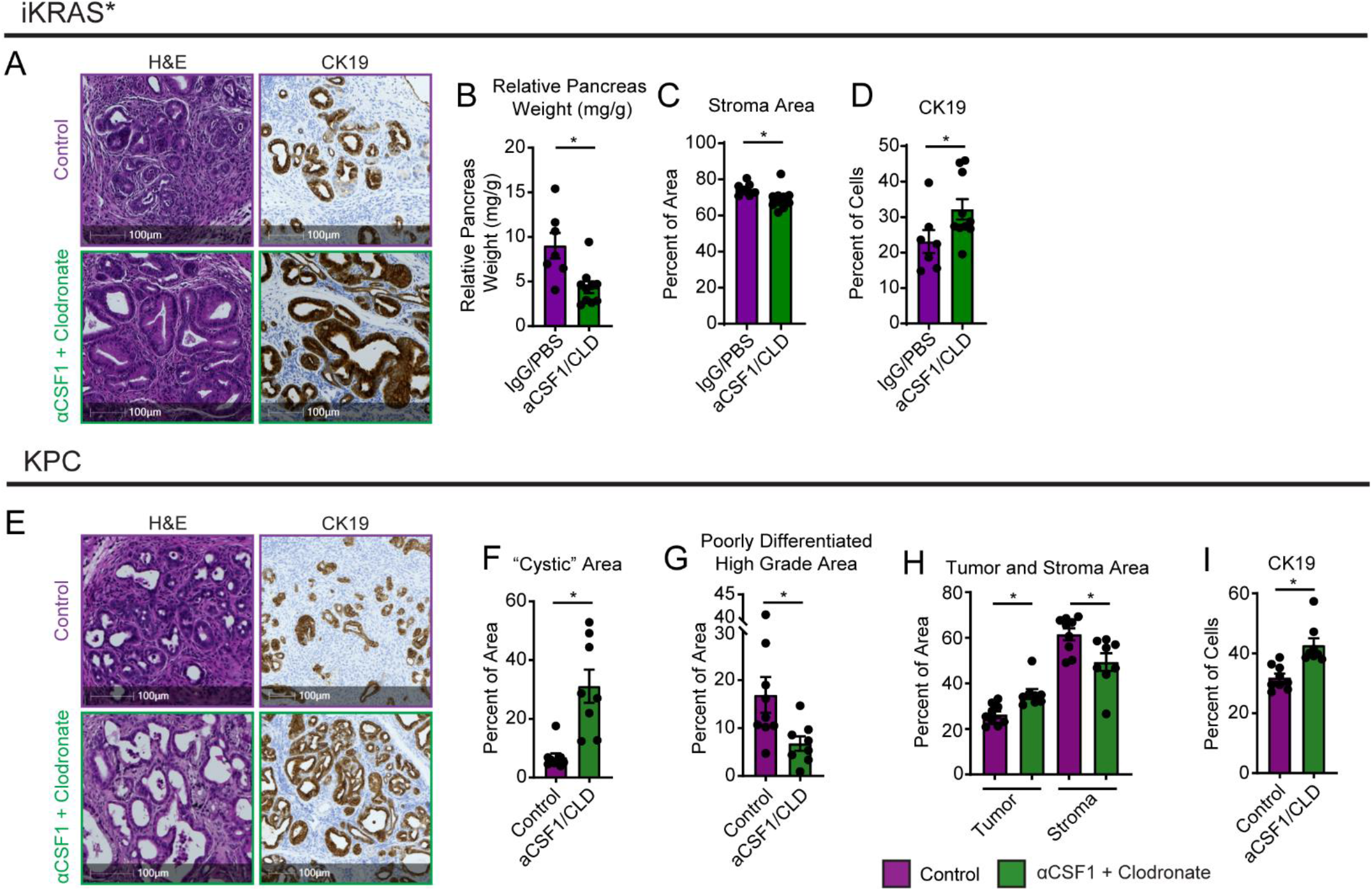
Tissue-resident macrophages drive fibrosis and pancreatitis-accelerated PDAC progression. **A**. Representative images of pancreas tissue stained for H&E and CK19 from p48-Cre;ROSA26-rtTa-IRES-EGFP;TetO-KRAS^G12D^ (iKRAS*) mice treated with IgG-PBS or αCSF1-CLD. **B**. Quantification of relative pancreas weight (mg pancreas tissue per g body weight); n = 7-9 mice/group. **C**. Quantification of the stromal area as percent of total pancreas area; n = 7-9 mice/group. **D**. Quantification of CK19 as the percentage of total cells; n = 7-9 mice/group. **E**. Representative images of pancreas tissue stained for H&E and CK19 from p48-Cre;LSL-KRAS^G12D^;p53^fl/+^ (KPC) mice treated with IgG-PBS or αCSF1-CLD. **F**. Quantification of tumor area presenting cyst-like appearance; n = 8-9 mice/group. **G**. Quantification of poorly differentiated/high grade invasive pancreatic tumor lesions as percent of total pancreas area, n = 8-9 mice/group. **H**. Quantification of tumor lesions and stromal areas as percent of total pancreas area; n = 8-9 mice/group. **I**. Quantification of CK19 as the percentage of total cells; n = 8-9 mice/group. Data are presented as mean ± SEM unless otherwise indicated. n.s., not significant; *p <0.05. For comparisons between two groups, Student’s two-tailed t-test was used.

Next, to corroborate these findings, we used a spontaneous PDAC GEMM, where pancreas-specific Cre expression causes activation of KRAS^G12D^ and single-copy loss of p53. These mice, abbreviated as KPC, spontaneously develop tumors by 6 months of age. To similarly study tumor progression as in the iKRAS* model, KPC mice were treated with αCSF1-CLD, given a recovery period, then briefly treated with cerulein to cause tissue damage and inflammation, and accelerate tumor growth (**Supp. Fig. 7b**). Pancreas tissue was then taken at 3.75 months of age, a timepoint when frank carcinomas can be observed. KPC tumors arising in TRM-depleted mice showed areas of well-differentiated tumor lesions with cyst-like appearance present with very little stroma (**Fig. 7f**). In contrast, quantitating poorly differentiated high grade invasive tumor lesions showed a 50% reduction in TRM-depleted mice compared to controls (**Fig. 7g**), suggesting TRMs are likely driving PDAC lesion progression. Consistent with TRM-depletion abrogating the stromal response, quantitation of stromal area showed a 10% reduction with αCSF1-CLD treatment, as well as an increased percentage of CK19^+^ tumor cells (**Fig. 7h-i**). Additionally, to confirm TRM depletion, F4/80 staining showed significant reduction in αCSF1-CLD treated mice (**Supp. Fig. 7c**). These KPC studies agree with the above iKRAS* data, and together suggest that in oncogenic settings, TRMs support tumor growth. As with pancreatitis, TRMs act by inducing fibrosis however instead of being beneficial for tissue protection and mouse survival, lead to promoting tumor growth and differentiation.

## Discussion

Tissue-resident macrophages (TRMs) are established during fetal development from embryonic progenitors. At the tissue-specific level, these cells undergo replacement from monocyte-derived sources to varying extents. Our findings here agree with previous studies of pancreas macrophages (Calderon *et al*., 2015; Zhu *et al*., 2017), showing that a significant portion of macrophages are long-lived TRMs established in part from embryonic progenitors. TRMs in the pancreas do not rely on continual replenishment from monocytes, and display distinct transcriptional phenotype and anatomical location from monocyte-derived macrophages (MDMs). Further, we identify a subset of TRMs that are primarily derived from embryonic progenitors and are marked as Lyve1^Hi^Cx3cr1^Lo^MHCII^Lo^. Other macrophage populations display mixed origin and are marked as Lyve1^Lo^Cx3cr1^Hi^MHCII^Hi^, agreeing with the dichotomy of marker expression by TRM populations that have been established across fat, dermis, heart, lung, mesenteric membranes, and other tissues (Chakarov *et al*., 2019; Dick *et al*., 2022; Gibbings et al., 2017; Lim *et al*., 2018; Zhang *et al*., 2021). Additionally, these other studies on Lyve1^Hi^MHCII^Lo^ TRMs in the lung, arteries, and mesenteric membranes report their involvement in extracellular matrix deposition and remodeling, and display a strikingly similar transcriptional profile as Lyve1^Hi^MHCII^Lo^ macrophages reported here in the pancreas.

It is not fully understood to what extent macrophage origin dictates its function, and how important tissue residence and a cell’s specific microenvironment are. Recent lineage-tracing studies have highlighted the ability of monocyte-derived cells to replace TRMs over time, and despite coming from a different origin, adopt similar marker expression and transcriptional phenotype (Liu *et al*., 2019). Additionally, transcriptional, and epigenetic profiling approaches have been used to show macrophages retain a remarkable amount of plasticity, and adoptively transferring even fully differentiated TRMs into a different tissue can cause reprogramming of up to 70% of genes (Lavin *et al*., 2014). Our study would agree with this, as scRNAseq data did not reveal completely distinct clustering or phenotype-based on origin, but rather that most macrophage clusters had varying contributions from both embryonic and HSC sources. Additionally, imaging of Lyve1^+^ TRMs revealed distinct anatomical location between lobules of the pancreas within the more highly fibrotic areas, suggesting these cells may be especially apt at coordinating fibrosis due to their specific microenvironment within the stroma.

Our study further suggests that the tissue remodeling program driven by TRMs leads to disparate outcomes depending on the type of tissue injury. TRM-driven fibrosis was critically important for mouse survival and recovery from pancreatitis through protective effects that limit acinar cell death and tissue damage by coordinating critical fibrotic responses. Indeed, it is thought that pancreatic stellate cell (PSC) activation is critical in the recovery process following inflammatory injury. PSCs have been shown to express growth factors and cytokines in vitro that may be critical for acinar cell regeneration (Omary *et al*., 2007), and histological analysis of human acute necrotizing pancreatitis specimens have shown close association and enhanced proliferation of PSCs and hypercellular regenerative spheres of ductal cells (Zimmermann, 2002). PSC activation for the wound healing process must, however, be transient. In the chronic setting, prolonged PSC activation has proven detrimental and a significant source of fibrosis that can interfere with pancreas function (Klöppel *et al*., 2003; Omary *et al*., 2007). Our studies here are somewhat limited to the acute setting able to be modeled in mice, where inflammation eventually resolves upon cerulein withdrawal. However, possible future directions include understanding how TRM-fibroblast interactions change under the chronic setting, and whether targeting these interactions could be used therapeutically to reduce fibrosis in chronic pancreatitis.

Similar to chronic pancreatitis, tumor conditions present prolonged inflammation and wound healing responses. Here, the same TRM-driven fibrotic response is co-opted by tumor cells to elicit a more favorable microenvironment and promote tumor growth. While still debated, multiple studies have shown that fibrosis can play a tumor supportive role, and some have even implicated macrophages in initiating or driving the fibrotic response (Chandler et al., 2019; DeNardo and Ruffell, 2019; Jiang et al., 2016; Zhang *et al*., 2021). Fibrotic pathologies have also been linked to a higher risk of developing cancer, such as chronic pancreatitis being a significant risk factor for PDAC (Lowenfels et al., 1993; Vujasinovic et al., 2020). In addition to TRMs driving fibrosis, other studies have implicated TRMs in supporting tumor growth through distinct mechanisms. In lung adenocarcinoma, TRMs are responsible for coordinating T regulatory cell responses, and when depleted, tumors grow more slowly (Casanova-Acebes *et al*., 2021). Similarly, in metastatic ovarian cancer models, TRMs have been shown to promote tumor cell growth and metastatic spreading (Etzerodt et al., 2020; Zhang *et al*., 2021). While the specific mechanisms by which TRMs promote tumor growth seem to vary depending on tissue and tumor model, there are some striking similarities. Most TRM populations display an alternative activation phenotype, secreting many growth factors and cytokines that either directly promote tumor cell growth, contribute to tumor-protective fibrosis, or disrupt anti-tumor T cell responses. Our findings here align with these previous studies, with Lyve1^Hi^MHCII^Lo^ TRMs displaying alternative activation, and expressing collagen and ECM remodeling proteins to contribute to fibrosis capable of protecting tumor cells and supporting their growth.

Aside from the functionality of TRMs, other studies have similarly focused on the roles of MDMs during pancreatitis, largely concluding that these cells are responsible for promoting inflammation and tissue damage (Saeki *et al*., 2012). Our findings here support these conclusions, as our transcriptional profiling revealed upregulation of inflammatory pathways in MDMs, but ultimately, we were not able to demonstrate functional outcomes when targeting MDMs. Depleting MDMs with CCR2-knockout mice displayed no changes in ADM, tissue damage, or mouse survival, perhaps due to CCR2-independent mechanisms by which inflammatory macrophages could accumulate in the pancreas during pancreatitis.

In summary, our study demonstrates that macrophage populations in the pancreas contain cells from both embryonic and monocyte sources. Upon tissue damage, TRMs drastically expand in number to primarily help coordinate a fibrotic response to protect the exocrine pancreas from further damage and cell death. While TRM-driven protective fibrosis is critically important for mouse survival and recovery from pancreatitis, it plays a tumor-supportive role during PDAC. Understanding the nuances of macrophage phenotypes and how they act during different pathologies, will be critical in developing therapies targeting these cells.

## Author Contributions

J.M.B. and D.G.D. conceived of and designed experiments, and wrote manuscript with input from all authors. J.M.B., C.Z., L.I.K., A.A.L., N.C.B., B.L.K., S.J.B., and D.G.D. performed experiments and analyzed data. N.C.B., M.A.L., N.Z., K.W.K., R.C.F., L.D., J.C.M., and G.J.R. provided key resources, expertise, input, and tissues.

## Acknowledgments

J.M.B. was funded by pre-doctoral fellowship F31 DK122633. D.G.D. and study costs were supported by NCI R01CA273190, R01CA177670, P50CA196510, P30CA09184215, and the BJC Cancer Frontier Fund. Collaborative studies with G.J.R. were supported by RO1AI049653. L.I.K. was funded by 5T32EB021955 2019-2021. K.W.K. was funded by NIDDK R01DK126753. We thank the Washington University Center for Cellular Imaging, the Flow Cytometry & Fluorescence Activated Cell Sorting Core and Genome Technology Access Center which are funded by D.G.D. fund P50CA196510.

## STAR Methods

### Resource Availability

#### Lead Contact

Further information and requests for resources and reagents should be directed to and will be fulfilled by the lead contact, David G. DeNardo.

#### Materials Availability

This study did not generate new unique reagents.

#### Data and Code Availability

This paper does not report original code, any information required to analyze the data reported is available from lead contact upon request.

### Experimental Model and Subject Details

#### Mouse Models

The following mouse models were purchased from Jackson Laboratories: CCR2^-/-^, Rosa26-LSL-eYFP, Csf1r-mer-iCre-mer. Lyve1-Cre;CSF1R^fl/fl^ (Lyve1^ΔCSF1R^) were a gift from Dr. Gwendalyn Randolph. LSL-tdTomato mice in the FvB background were a gift from Dr. Gregory Longmore, and were crossed to Csf1r-mer-iCre-mer for lineage tracing studies. Flt3-Cre mice were a gift from Dr. Thomas Boehm, and were crossed to Rosa26-LSL-eYFP. For Flt3-Cre;Rose26-LSL-eYFP experiments, male mice were used between ages of 8-16 weeks. All other experiments were performed on both male and female mice between the ages of 8-16 weeks. Mice were maintained in the Laboratory for Animal Care barrier facility at the Washington University School of Medicine. Washington University School of Medicine Institutional Animal Studies Committee approved all animal studies.

#### Murine Tumor Models

KPC mice (p48-Cre;LSL-KRAS^G12D^;p53^fl/+^) used in these studies have been previously described. They were backcrossed to C57BL/6 mice and tested for congenic markers. iKRAS* mice (Pft1a-Cre;LSL-rtTa;TetO-KRAS^G12D^) were a kind gift from Dr. Marina Pasca di Magliano, and have been previously characterized. Briefly, Kras^G12D^ expression was induced by administration of doxycycline in drinking water at a concentration of 0.5mg/mL, supplemented with 3% sucrose w/v. Upon start of doxycycline, mice were treated with acute cerulein on two consecutive days, each day injecting 100ug/kg of cerulein every hour for 6 hours. Doxycycline administration was continued for 3 weeks, replacing drinking water every 3 days. KPC-1 cell line was derived from a 2.2 month old KPPC mouse (p48-Cre;LSL-KRAS^G12D^;p53^fl/fl^), and grown on collagen-coated tissue culture flasks for < 12 generations. To establish orthotopic pancreas KPC tumor models, Flt3-YFP or C57BL/6 mice were implanted with 100,000 KPC-1 cells in 5uL serum-free DME/F-12 and 45uL Cultrex (Trevigen) by injecting cells directly into the pancreas.

For lung tumors, the KPL86 cell line was derived from lung tissues of the 9-month-old *Kras*^*LSL-G12D*^*/p53*^*flox/+*^ mouse treated with adenovirus-delivered Cre-recombinase. Cells were grown on collagen-coated tissue culture flasks for < 10 passages. To establish orthotopic NSCLC models, 1,000,000 KPL86 cells in 100 μL of phosphate-buffered saline (PBS, Trevigen, Gaithersburg, MD, USA) were retro-orbitally injected into 8–12-week-old Flt3-YFP mice.

For liver metastasis of PDAC cells, hemispleen injection of 500,000 KPC-2 (aka “KP2”) tumor cells was performed using an adapted procedure from published studies (Lee et al., 2019; Soares et al., 2014). The KP2 cell line was derived from pancreatic tumors of a 6-month-old p48-CRE+/LSL-KRAS^G12D^/p53^flox/+^ mouse, and tumor cells additionally were modified to express mCherry and click beetle red luciferase. To implant cells in the liver, the mice were anesthetized under continuously-monitored isoflurane, and the spleen was exteriorized from the peritoneal cavity under sterile conditions through a left flank laparotomy. The spleen was ligated using medium titanium Horizon ligation clips (Teleflex, Morrisville, NC) and divided in half. Tumor single-cell suspension in sterile PBS (100 uL), followed by 150uL of sterile PBS flush, was injected into the inferior pole of the spleen with a 30-gauge needle (BD, Franklin Lakes, NJ), monitoring for blanching of the spleen parenchyma and splenic vein to ensure injection efficacy. Once the cells and PBS flush were injected, the splenic vein was ligated, and the inferior pole of the spleen was excised. Once hemostasis was ensured, the peritoneum was closed with 5-0 ligature (Ethicon, Raritan, NJ), and the skin was closed with the AutoClip system (Braintree Scientific, Braintree, MA). Mice were post-operatively monitored and managed per IACUC guidelines for survival surgery. Livers were harvested at day 14 for cell sorting and single cell RNA sequencing analysis.

#### Human samples

Human PDAC (with adjacent non-tumor “normal” pancreatic tissue) and pancreatitis (with adjacent “normal” pancreatic tissue) samples were obtained from consenting patients diagnosed at Barnes-Jewish Hospital. The studies described were approved by the Washington University Institutional Review Board under protocol numbers 201704078 and 201908148. Adjacent normal pancreas was identified by pathologist by H&E-stained slides.

### Method Details

#### Experimental Pancreatitis

Acute pancreatitis was induced in mice by administering 6 hourly intraperitoneal injections (i.e. once per hour for six hours) of cerulein at a dose of 100ug/kg given every other day for one week.

For severe acute pancreatitis via osmotic pump delivery of cerulein, osmotic pumps (Alzet Osmotic Pumps, Cupertino, CA) were loaded with 100uL of concentrated cerulein (8ug cerulein per day was loaded with 1.33mg/mL and 10ug cerulein per day was loaded with 1.67mg/mL). Pumps were then implanted into the peritoneal cavity of mice, and body weight was monitored every day for up to 14 days.

#### Macrophage Depletion

Tissue-resident macrophages were depleted with a combination of CSF1 neutralizing antibodies and clodronate-loaded liposomes. 8-12 week old FvB mice were treated with two doses of CSF1 antibody (BioXcell, 5A1, 1mg and 0.5mg on days -14 and -11 respectively), followed by two doses of clodronate-loaded liposomes (Liposoma, 200uL each dose) on days -13 and -10. Mice were then allowed to recover for 10 days to allow monocytes and monocyte-derived macrophages to reconstitute. On day 0, mice were either treated with cerulein by IP injection (described above) or implanted with osmotic pump to deliver cerulein.

Similarly, KPC mice were treated with two doses of αCSF1 (1mg and 0.5mg) separated by two days and starting at 10 weeks of age. Subsequent days after αCSF1 treatment, mice were injected with 100uL of clodronate-loaded liposomes. Mice were then allowed to recover for ∼2 weeks, then given brief cerulein treatment starting at 12 weeks of age by giving 6 hourly injections of 100ug/kg cerulein every other day for 5 days. Following cerulein, tumors were allowed to grow for ∼3 weeks and mice were sacrificed at 15 weeks of age.

iKRAS* mice were treated with a similar regimen of two doses of αCSF1 (1mg and 0.5mg) 14 and 11 days before starting doxycycline administration. Following αCSF1 treatment, mice were injected with clodronate-loaded liposomes (100uL) on subsequent days. After αCSF1 and clodronate treatment, mice were allowed to recover for 10 days, after which doxycycline was given in drinking water (0.5mg/mL with 3% sucrose replaced every 3 days, as above) for ∼3 weeks. Additionally, on days 0 and 1, iKRAS* mice were treated with acute cerulein (100ug/kg) by 6 hourly injections.

#### Lineage Tracing

For lineage tracing of embryonically derived macrophages, timed breeding pairs were set up by crossing Csf1r-mer-iCre-mer mice with Rosa26-LSL-tdTomato mice (both on FvB background). Embryonic timeline was estimated by observation of vaginal plug, with 12 pm on the day of plug formation being 0.5 days post coitum (dpc). Pregnant mice were then treated with tamoxifen (75ug/g) supplemented with progesterone (37.5ug/g) dissolved in corn oil at ∼9.5dpc. Offspring of tamoxifen-pulsed mice were then treated with vehicle or acute cerulein (as described above) at 6 weeks of age and sacrificed for fate mapping analyses.

For lineage tracing of resident macrophages, adult Cx3cr1-CreERT2;LSL-tdTomato and Csf1r-mer-iCre-mer;LSL-tdTomato mice were pulsed with tamoxifen (75mg/kg) dissolved in corn oil on five consecutive days. Mice were then allowed a chase period or 3-10 weeks, or until blood monocytes lost tdTomato labeling. Following chase period, mice were treated with vehicle or acute cerulein as described above, then sacrificed for flow cytometry or immunohistochemistry analyses.

#### Mouse Tissue Isolation and Flow Cytometry

Mice were sacrificed by cardiac perfusion with ∼10-15mL PBS-heparin under isofluorane anesthesia. Pancreas or tumor tissue was then isolated, minced with scissors, and digested in ∼15-25mL Dulbecco’s Modified Eagle Medium (DMEM) with 2mg/mL Collagenase A and 1x DNase I for either 15 minutes (steady-state pancreas and cerulein treated pancreas) or 25 minutes (tumor tissue) at 37°C with constant stirring. Digestion buffer was quenched with 3mL fetal bovine serum (FBS) and filtered through 100uM Nylon mesh, pelleted by centrifugation (2000rmp for 5 min) and resuspended in FACS buffer (1% BSA in PBS). Brain tissue was minced and triturated through 40uM Nylon mesh and resuspended in FACS buffer to obtain single-cell suspension.

Cell suspensions were then blocked with rat anti-mouse CD16/CD32 antibodies for 10 minutes on ice, pelleted by centrifugation, and stained with fluorophore-conjugated extracellular antibodies for 25 minutes on ice. For samples with endogenous fluorescent protein labeling, cells were then washed twice with FACS Buffer, then run on BD Fortessa X20 immediately. For unlabeled cells, samples were washed twice following antibody staining, fixed with BD Fixation Buffer for 30 minutes on ice, washed, and resuspended in FACS Buffer. Fixed samples were stored at 4°C and run on BD Fortessa X20 within two weeks. For samples requiring intracellular staining, cells were washed following extracellular staining, and stained using eBioscience FoxP3 Transcription Factor Staining Kit according to instructions by manufacturer.

To quantify proliferating cells in Flt3-YFP mice, tissue was digested and extracellular staining was conducted as described above. Cells were then fixed in 4% paraformaldehyde for 10 minutes on ice, then permeabilized in ice-cold 70% ethanol for 3 hours, and stained with Ki-67 antibody diluted in FACS Buffer for 20 minutes on ice. Cells were then washed twice and immediately run on BD Fortessa X20.

To quantitate blood monocytes, 100uL blood was obtained by cardiac puncture prior to perfusions and deposited into PBS-heparin. Blood was then pelleted by centrifugation, resuspended in 5mL red blood cell lysis buffer for 10 minutes at room temperature, cells pelleted by centrifugation, then stained with fluorophore-conjugated antibodies for 25 minutes on ice. Cells were then fixed in BD Fixation Buffer for 30 minutes on ice, washed twice with FACS Buffer, and stored at 4°C and run on BD Fortessa X20 within two weeks. For experiments requiring blood quantitation prior to mice being sacrificed, blood was obtained by tail vein bleeding, resuspended in RBS Lysis Buffer, stained with extracellular antibodies as described above.

#### Fluorescence-Activated Cell Sorting (FACS)

Single-cell suspensions were obtained as described above. Cells were then resuspended in FACS Buffer containing CD16/CD32 antibodies to block for 10 minutes on ice, pelleted by centrifugation, then resuspended in fluorophore-conjugated antibodies, stained for 25 minutes on ice, then washed with FACS Buffer. Cells were then immediately sorted on Aria-II (BD Biosciences).

For homeostatic pancreas, pancreatitis, and pancreatic tumors, live macrophages (CD45+CD11b+CD3-CD19-B220-SiglecF-Ly6G-Ly6C-7AAD-F4/80+MHCII^HI/Lo^) were sorted. For Bulk RNAseq analyses, Flt3-YFP positive and negative macrophages were sorted directly into RNA Lysis Buffer (Omega Biotek) and RNA isolated using EZNA Kit (Omega Biotek). For single-cell RNA-sequencing (scRNAseq), cells were sorted into FACS buffer and library preparation was conducted as described below.

Liver single-cell suspensions were obtained as described above. Cells were then resuspended in anti-CD45 magnetic beads (BD Biosciences), incubated for 15 minutes on ice, washed in FACS buffer, and applied to LS magnetic column to enrich for CD45+ cells. Following magnetic isolation, cells were spun down, resuspended in fluorophore-conjugated antibodies, stained for 25 minutes on ice, washed with FACS buffer, then immediately sorted. Flt3-YFP positive and negative liver macrophages were sorted (CD45+CD11b+CD3-CD19-B220-SiglecF-Ly6G-7AAD-) into FACS buffer, then scRNAseq library preparation was conducted as described below.

Lung single-cell suspensions were obtained as described above. Cells were then resuspended in FACS Buffer containing CD16/CD32 antibodies to block for 10 minutes on ice, pelleted by centrifugation, then resuspended in fluorophore-conjugated antibodies, stained for 25 minutes on ice, then washed with FACS Buffer. Cells were then immediately sorted on Aria-II (BD Biosciences). Flt3-YFP positive and negative macrophages were sorted (CD45+CD3-CD19-B220-7AAD-CD11b+CD11c^Hi/Lo^F4/80+) into FACS buffer, then scRNAseq library preparation was conducted as described below.

For fibroblasts, homeostatic or cerulein treated pancreas tissue was taken, and single-cell suspensions were isolated as described above. Cells were then resuspended in FACS Buffer containing CD16/CD32 antibodies to block for 10 minutes on ice, pelleted by centrifugation, then resuspended in fluorophore-conjugated antibodies, stained for 25 minutes on ice, then washed with FACS Buffer. Cells were then immediately sorted on Aria-II (BD Biosciences). Total fibroblasts were sorted (CD45-EpCAM-CD31-PDPN+) into FACS Buffer, then scRNAseq library preparation was conducted as described below.

After cells were sorted into FACS Buffer, they were pelleted by centrifugation, resuspended in 0.04% BSA in PBS, and cell counts were obtained. Single-cell RNA-sequencing library preparation was then conducted by the Genome Technology Access Center at Washington University. Briefly, cells from each sample were encapsulated into droplets, and libraries were prepared using Chromium Single Cell 3’v3 Reagent kits according to the manufacturer’s protocol (10x Genomics, Pleasanton, CA, USA). The generated libraries were sequenced by a NovaSeq 6000 sequencing system (Illumina, San Diego, CA, USA) to an average of 50,000 mean reads per cell. Cellranger mkfastq pipeline (10X Genomics) was used to demultiplex illumine base call files to FASTQ files. Afterward, fastq files from each sample were processed with Cellranger counts and aligned to the mouse mm10 reference genome (Cellranger v.4.0.0, 10X Genomics, mouse reference mm10-2020-A from https://cf.10xgenomics.com/supp/cell-exp/refdata-gex-mm10-2020-A.tar.gz) to generate feature barcode matrix.

#### Mouse Bulk RNAseq Analysis

Pancreas macrophages were sorted as described above, directly into TRK Lysis buffer (Omega Bio-Tek). RNA was isolated by E.Z.N.A. Total RNA isolation kit per manufacturer’s recommendations (Omega Bio-Tek). Following quality control by bioanalyzer, libraries were prepared using the Clontech SMARTer kit (Takara Bio) at the Washington University Genome Technology Access Center (GTAC). Libraries were then indexed, pooled, and sequenced on Illumina HiSeq 3000 (Illumina). Illumina’s bcl2fastq software was then used for basecalls and demultiplexing. Reads were then aligned to the Ensembl release 76 top-level assembly with STAR version 2.0.4b, and gene counts were derived from the number of uniquely aligned unambiguous reads by Subread:featureCount version1.4.5. Gene counts were then imported into Edge,R and TMM normalization size factors were calculated to adjust for differences in library size. Limma and voomWithQualityWeights was then used to calculate weighted likelihoods based on the observed mean-variance relationship of every gene and sample. Differential expression analysis was then performed to analyze for differences between Flt3-YFP lineages and results filtered by Benjamini-Hochberg false discovery rate adjusted p-values less than or equal to 0.05. Global perturbations in known Gene Ontology (GO) terms and KEGG pathways were detected using GAGE to test for changes in expression of reported log 2 fold-changes reported by Limma in each term versus background log 2 fold-changes of all genes outside the respective term. Differentially expressed genes and gene sets were displayed using pHeatmap or the Phantasus online tool (Artyomov, https://artyomovlab.wustl.edu/phantasus/).

#### Mouse Single-cell RNAseq Analysis

The filtered feature barcode matrix from each sample was then loaded into Seurat as Seurat objects (Seurat v.3). For each Seurat object, genes that were expressed in less than three cells and cells that expressed less than 1,000 or more than 8,000 genes, were excluded. Cells with greater than 10% mitochondrial RNA content were also excluded, resulting in between 2297 and 17880 cells, as indicated in **Supplementary Table 1**. SCTransform with default parameters was used on each individual sample to normalize and scale the expression matrix against the sequence depths and percentages of mitochondrial genes (Hafemeister and Satija, 2019). Principle component analysis (PCA) was performed (function RunPCA). A UMAP dimensional reduction was performed on the scaled matrix using the first 25 PCA components to obtain a two-dimensional representation of cell states. Then, these defined 25 dimensionalities were used to refine the edge weights between any two cells based on Jaccard similarity (FindNeighbors), and were used to cluster cells through FindClusters functions.

To characterize clusters, the FindAllMarkers function with logfold threshold = 0.25 and minimum 0.25-fold difference and MAST test were used to identify signatures alone with each cluster. For macrophage/monocyte samples, the macrophage/monocytes clusters were selected, and the top 3,000 variable features were recalculated to recluster to a higher resolution. Macrophages were selected based on clusters with high expressions of known macrophage marker genes, including *Csf1r, C1qa, and C1qb*, and confirmed by the absence of *Cd3e, Ms4a1, Krt19, Zbtb46*, and *Flt3*, and further confirmed by identifying DEGs associated with potential macrophage clusters, when compared to known macrophage specific marker genes. For fibroblast samples, the fibroblast clusters were selected and similarly reclustered. Fibroblasts were selected based on clusters with high expression of known fibroblast marker genes, including *Pdpn, Pdgfra, Col1a1*, and confirmed by the absence of *Krt19, Prss2, Amy1a*. For GSEA comparisons, the log_2_ (fold-change) of all genes detected with min.pct > 0.1 and past MAST test was used as a ranking metric. GSEA was performed using GO terms, KEGG pathways, Reactome, and MSigDB gene sets with Benjamini-Hochberg FDR < 0.05 in ClusterProfiler (Wu et al., 2021). For DEGs between the two groups in each library, we filtered genes with a Bonferroni-corrected p-value < 0.05 and fold-change >1.2 or <0.8.

#### Human Single-cell RNAseq Analysis

For the human dataset (Zhou *et al*., 2021), cells with greater than 15% mitochondrial genes were excluded and cells that expressed less than 500 genes were excluded. SCTransform with default parameters was used on each individual sample to normalize and scale the expression matrix against sequence depth and percentage of mitochondrial genes (Hafemeister and Satija, 2019). Cell cycle scores and corresponding cell cycle phases were then calculated and assigned after SCTransform based on S and G2/M genes (CellCycleScoring). Differences between the S phase and G2/M phase were then regressed out by SCTransform on individual samples. Variable features were calculated for every sample in the dataset independently and randed based on the number of samples they were independently identified (SelectIntegrationFeatures). The top 3,000 variable features were then used for PCA (RunPCA). The calculated PCA embedding of each cell was then used as an input for the soft k-means clustering algorithm. Briefly, through iteration, the algorithm designated the cluster-specific centroids and cell-specific correction factors corresponding to batch effects. The correction factors were used to assign cells into clusters until the assignment was stable (RunHarmony). The first 20 PCA components were then used to refine the edge weights between any two cells based on Jaccard similarity (FindNeighbors), and were used to cluster cells through FindClusters function at a resolution of 0.3, resulting in 24 clusters. To characterize clusters, the FindAllMarkers function with logfold threshold = 0.25 and minimum 0.25-fold difference and MAST test were used to identify signatures alone with each cluster. Macrophage/monocyte clusters were then selected, and the top 3,000 variable features were recalculated to recluster to a higher resolution, resulting in 5970 total cells and 8 clusters. Macrophages were selected based on clusters with high expressions of known macrophage marker genes, including *CSF1R, C1QA, and C1QB, CD68*, and confirmed by the absence of *CD3E, MS4A1, KRT19, CD1C*, and further confirmed by identifying DEGs associated with potential macrophage clusters, when compared to known macrophage specific marker genes.

#### Immunohistochemistry and Immunofluorescence

Mouse tissues were fixed in 10% formalin overnight at 4°C, dehydrated through graded ethanol washes of 30% ethanol, 50% ethanol, and 70% ethanol for 20 minutes each. Tissue was then run through Leica ASP 6025 tissue processor and embedded in paraffin, then sectioned at 6-μm thick tissue sections. Immunohistochemistry staining was then performed on Bond Rxm autostainer (Leica Biosystems) according to manufacturer’s recommendations. Briefly, paraffin slides were loaded onto machine, baked at 60°C for 15 minutes, dewaxed, rehydrated, antigens were retrieved with either citrate or EDTA antigen retrieval solution (Leica Biosystems). Endogenous peroxidase activity was blocked with a peroxide block, primary antibodies were then applied for 60 minutes. Either species-specific or biotinylated secondary antibodies were then applied to the slides. Next, either DAB or Fast Red chromogens were applied, finally followed by hematoxylin counterstain. Slides stained with DAB were then dehydrated through graded ethanols, xylene, then mounted with Cytoseal XYL (Thermo Fisher), and slides stained with Fast Red were air dried for 15 minutes then mounted with Vectamount Permanent Mounting Medium (Vector Laboratories).

#### Multiplex Immunohistochemistry Staining

Formalin-fixed paraffin-embedded (FFPE) tissue was sectioned at 6-μm, loaded onto Bond Rxm autostainer, baked for 60 minutes at 60°C, dewaxed and rehydrated. Series of staining was performed with multiple markers as indicated, based on previously published study (Tsujikawa et al., 2017). Briefly, slides were first blocked with peroxide block, goat serum block, and species-specific F’ab block (Jackson Laboratories), then primary antibodies were incubated for 60 minutes, followed by species-specific secondaries conjugated to horseradish peroxidase (HRP). For primary antibodies raised in rabbit, Polymer (anti-rabbit poly-HRP, Leica Biosystems) was used as a secondary antibody, for primary antibodies raised in mouse, Post-Primary (rabbit anti-mouse, Leica Biosystems) secondary was used, followed by Polymer, for other primary antibodies, biotinylated species-specific secondaries were used, followed by streptavidin-HRP (SA-HRP, Leica Biosystems)(anti-rabbit poly-HRP and SA-HRP are provided in Intense R and Polymer Refine Detection Kits respectively, Leica Biosystems). Slides were then chromogenically visualized with AEC (Invitrogen) and counterstained with hematoxylin (Dako). Stained slides were then mounted with Vectamount Aqueous Mounting Medium (Vector Laboratories) and imaged on Zeiss Axio Scan.Z1 (Zeiss) at 20X magnification. Before the next round of staining, coverslips and mounting medium was removed by soaking slides in TBST (1X TBS with 0.05% Tween-20) for several hours, then destained by graded ethanols (50% ethanol, 70% ethanol with 1% HCl, 100% ethanol, 70% ethanol, 50% ethanol, and DI water for 5-10 minutes each) then loaded onto Bond Rxm for staining of next marker.

### Quantification and Statistical Analysis

Number of mice and statistical tests used are reported in each figure legend. Statistical analyses were performed by Graphpad Prism v9, using unpaired Student’s t-test, ANOVA analysis (Bonferroni multiple comparisons), or unpaired non-parametric Mann-Whitney U test as appropriate for normality of data. Log-Rank (Mantel-Cox) test was used for survival experiments. Data displayed are mean ± SEM, unless otherwise noted. p<0.05 is considered statistically significant for all studies; *p<0.05, n.s. denotes not significant.

## Key Resources Table

| REAGENT or RESOURCE | SOURCE | IDENTIFIER |
| --- | --- | --- |
| **Antibodies** |  |  |
| Anti-mouse B220-APC (Clone RA3-6B2) | BioLegend | Cat# 103212 |
| Anti-mouse CD11b-Alexa700 (Clone M1/70) | eBioscience | Cat# 56-0112-82 |
| Anti-mouse CD16/CD32 (Clone 93) | eBioscience | Cat# 14-0161-82 |
| Anti-mouse CD19-APC (Clone eBio1D3) | eBioscience | Cat# 17-0193-82 |
| Anti-mouse CD45-APC-eFluor780 (Clone 30-F11) | eBioscience | Cat# 47-0451-82 |
| Anti-mouse CD45-PE-Cy7 (Clone 30-F11) | eBioscience | Cat# 25-0451-82 |
| Anti-mouse CD115-BV605 (Clone AFS98) | BioLegend | Cat# 135517 |
| Anti-mouse CD115-PE (Clone AFS98) | eBioscience | Cat# 12-1152-82 |
| Anti-mouse CX3CR1-PE (Clone SA011F11) | BioLegend | Cat# 149006 |
| Anti-mouse F4/80-PE (Clone BM8) | eBioscience | Cat# 12-4801-82 |
| Anti-mouse F4/80-PE-Cy5 (Clone BM8) | eBioscience | Cat# 15-4801-82 |
| Anti-mouse Ki-67-BV605 (Clone 16A8) | BioLegend | Cat# 652413 |
| Anti-mouse Ki-67-FITC (Clone SolA15) | eBioscience | Cat# 11-5698-82 |
| Anti-mouse Ly6C-Alexa488 (Clone HK1.4) | eBioscience | Cat# 53-5932-82 |
| Anti-mouse Ly6C-PerCP-Cy5.5 (Clone HK1.4) | eBioscience | Cat# 45-5932-82 |
| Anti-mouse Ly6G-APC (Clone 1A8) | BioLegend | Cat# 127614 |
| Anti-mouse Ly6G-PE (Clone 1A8) | eBioscience | Cat# 12-9668-82 |
| Anti-mouse MHCII-APC-eFluor780 (Clone M5/114.15.2) | eBioscience | Cat# 47-5321-82 |
| Anti-mouse MHCII-eFluor450 (Clone M5/114.15.2) | eBioscience | Cat# 48-5321-82 |
| Anti-mouse SiglecF-APC (Clone ES22-10D8) | Miltenyi | Cat# 130-102-241 |
| Anti-mouse Amylase (Polyclonal) | Millipore Sigma | A8273 |
| Anti-mouse Keratin 19 (Clone TROMA-III) | University of Iowa DSHB | TROMA-III |
| Anti-mouse Sox9 (Polyclonal) | Millipore Sigma | ABE571 |
| Anti-mouse Ki-67 (Clone D3B5) | Cell Signaling | 12202 |
| Anti-mouse Fibronectin (Polyclonal) | Abcam | Ab23750 |
| Anti-mouse Podoplanin (Clone RTD4E10) | Abcam | Ab11936 |
| Anti-mouse Type 1 Collagen (Polyclonal) | Southern Biotech | 1310-01 |
| Anti-mouse F4/80 (Clone D2S9R) | Cell Signaling | 70076 |
| Anti-mouse Lyve1 (Clone Aly7, biotinylated) | eBioscience | 13-0443-82 |
| Anti-RFP (Polyclonal) | Rockland | 600-401-379 |
| Anti-GFP (Clone EPR14104) | Abcam | Ab183734 |
| Anti-Keratin 17/19 (Clone D4G2) | Cell Signaling | 12434 |
| Anti-Cleaved Caspase 3 (Clone ASP175) | Cell Signaling | 9661S |
| Anti-Rat Rabbit Secondary | Vector | AI-4001 |
| Anti-Rat Biotin Secondary | Vector | BA-9400 |
| Anti-Syrian Hamster Biotin | Abcam | Ab7145 |
| Chemicals, peptides, and recombinant proteins |  |  |
| Anti-Rabbit Fab | Jackson Immuno Research | 111-007-0039 |
| Anti-Rat Fab | Jackson Immuno Research | 712-007-003 |
| Cytoseal XYL | Thermo Fisher | 8312-4 |
| Vectamount Permanent Mounting Medium | Vector | H-5000 |
| Vectamount Aqueous Mounting Medium | Vector | H-5501 |
| CSF1 neutralizing antibody (Clone 5A1) | BioXCell | BE0204 |
| PBS Liposomes & Clodronate Liposomes | Liposoma | CP-005-005 |
| Tamoxifen | Sigma-Aldrich | T5648 |
| Progesterone | Sigma-Aldrich | P0130 |
| CD45 MicroBeads | Miltenyi Biotec | 130-052-301 |
| LS Columns | Miltenyi Biotec | 130-042-401 |
| Critical commercial assays |  |  |
| Polymer Refine Detection Kit | Leica Biosystems | DS9800 |
| Polymer Refine RED Detection Kit | Leica Biosystems | DS9390 |
| Intense R Detection Kit | Leica Biosystems | DS9263 |
| AEC Substrate Kit | Abcam | Ab64252 |
| Dual Endogenous Enzyme Block | Dako | S2003 |
| Avidin/Biotin Blocking Kit | Vector | SP-2001 |
| E.Z.N.A. Total RNA Kit I | Omega | R6834-02 |
| Trichrome Kit | Diagnostic Biosystems | Cat#KT034 |
| Experimental models: Cell lines |  |  |
| KPC-1 | (Mitchem et al., 2013) | N/A |
| KPL86 |  |  |
| KPC-2-mCherry | (Jiang *et al.*, 2016) | N/A |
| Experimental models: Organisms/strains |  |  |
| C57BL/6J mice | Jackson Laboratory | Cat#000664 |
| FVB/NJ mice | Jackson Laboratory | Cat#001800 |
| CCR2 knockout mice | Jackson Laboratory | Cat# 004999 |
| CSF1R-mer-iCre-mer mice | Jackson Laboratory | Cat# 019098 |
| Rosa26-LSL-tdTomato mice | Laboratory of Gregory Longmore | N/A |
| Flt3-Cre mice | Laboratory of Thomas Boehm | N/A |
| Rosa26-eYFP mice | Jackson Laboratory | Cat#006148 |
| CX3CR1-CreERT2 mice | Jackson Laboratory | Cat#025524 |
| CX3CR1-GFP mice | Jackson Laboratory | Cat#005582 |
| Lyve1-Cre mice | Laboratory of Gwendalyn Randolph | N/A |
| CSF1R-flox mice | Laboratory of Gwendalyn Randolph | N/A |
| p48-Cre;ROSA26-rtTa-IRES-EGFP;TetO-KRAS<sup>G12D</sup> (iKRAS*) mice | Laboratory of Marina pasca di Magliano | N/A |
| p48-Cre mice | Laboratory of Sunil Hingorani | N/A |
| LSL-Kras<sup>G12D</sup> mice | Jackson Laboratory | Cat#008179 |
| p53<sup>fl/fl</sup> mice | Jackson Laboratory | Cat#008462 |
| Software and algorithms |  |  |
| Flowjo v10.7.2 | Flowjo, L.L.C. | Flowjo, L.L.C. |
| Prism v9 | Graphpad | www.graphpad.com |
| HALO v3.2.1851.393 | Indica Labs | https://www.indicalab.com |
| FACSDiva | BD Biosciences | RRID: SCR_001456 |
| R v3.6.3 | R Core Team | https://cran.r-project.org/bin/windows/base/old/3.6.3/ |
| Clusterprofiler | https://github.com/YuLab-SMU/clusterProfiler |  |
| Seurat v 3.2.0 | https://satijalab.org/seurat/ |  |
| Harmony | https://github.com/immunogenomics/harmony |  |

