## Supplemental Figures for "Pancreas resident macrophage-induced fibrosis has divergent roles in pancreas inflammatory injury and PDAC"

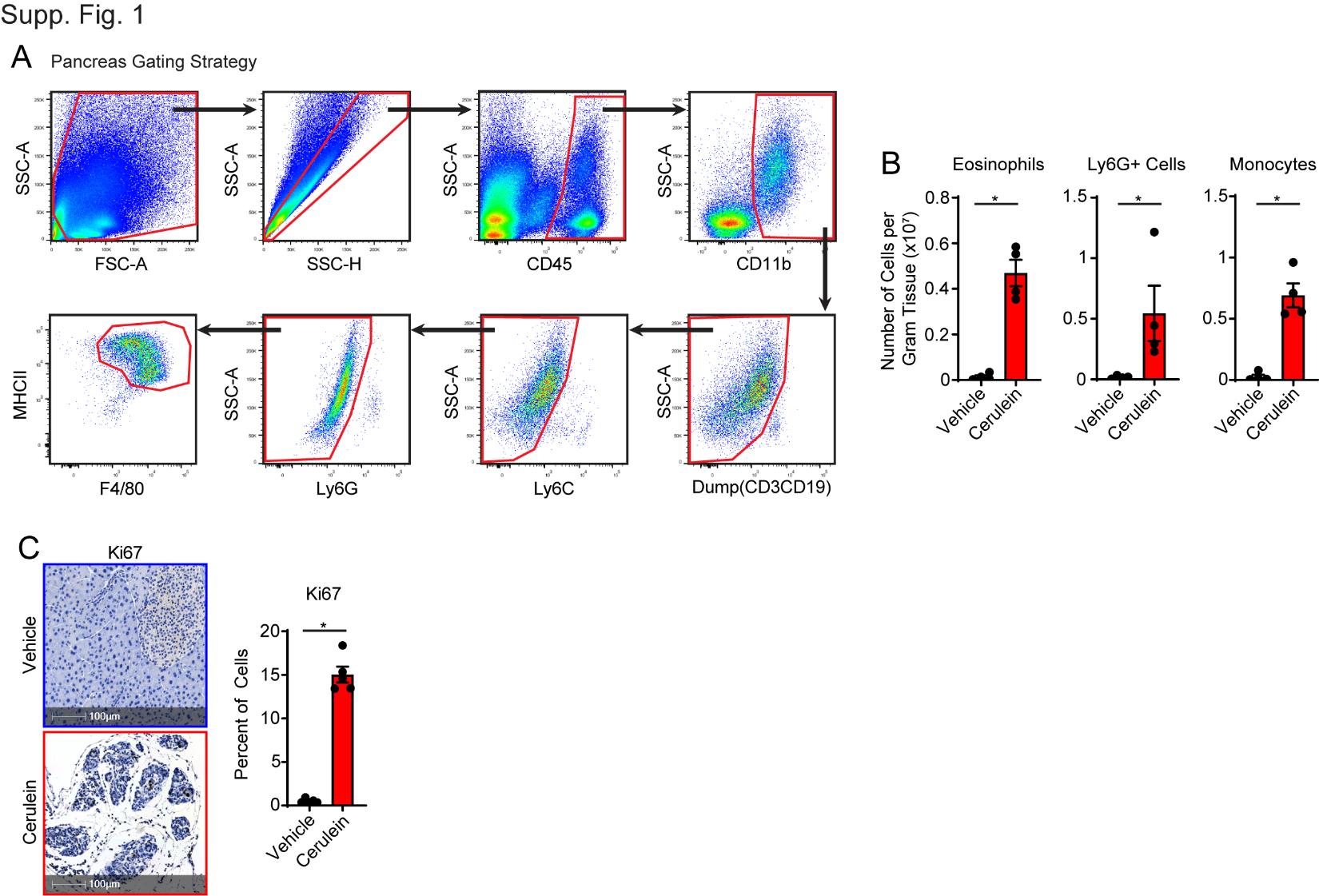

**Supplemental Figure 1: A.** Gating strategy for pancreas macrophages. **B.** Density of eosinophils, granulocytes, and monocytes in pancreas of vehicle and cerulein treated mice; n = 4 mice/group. **C.** Immunohistochemistry and quantification of Ki67 in pancreas tissue of vehicle and cerulein treated mice; n = 4 mice/group.

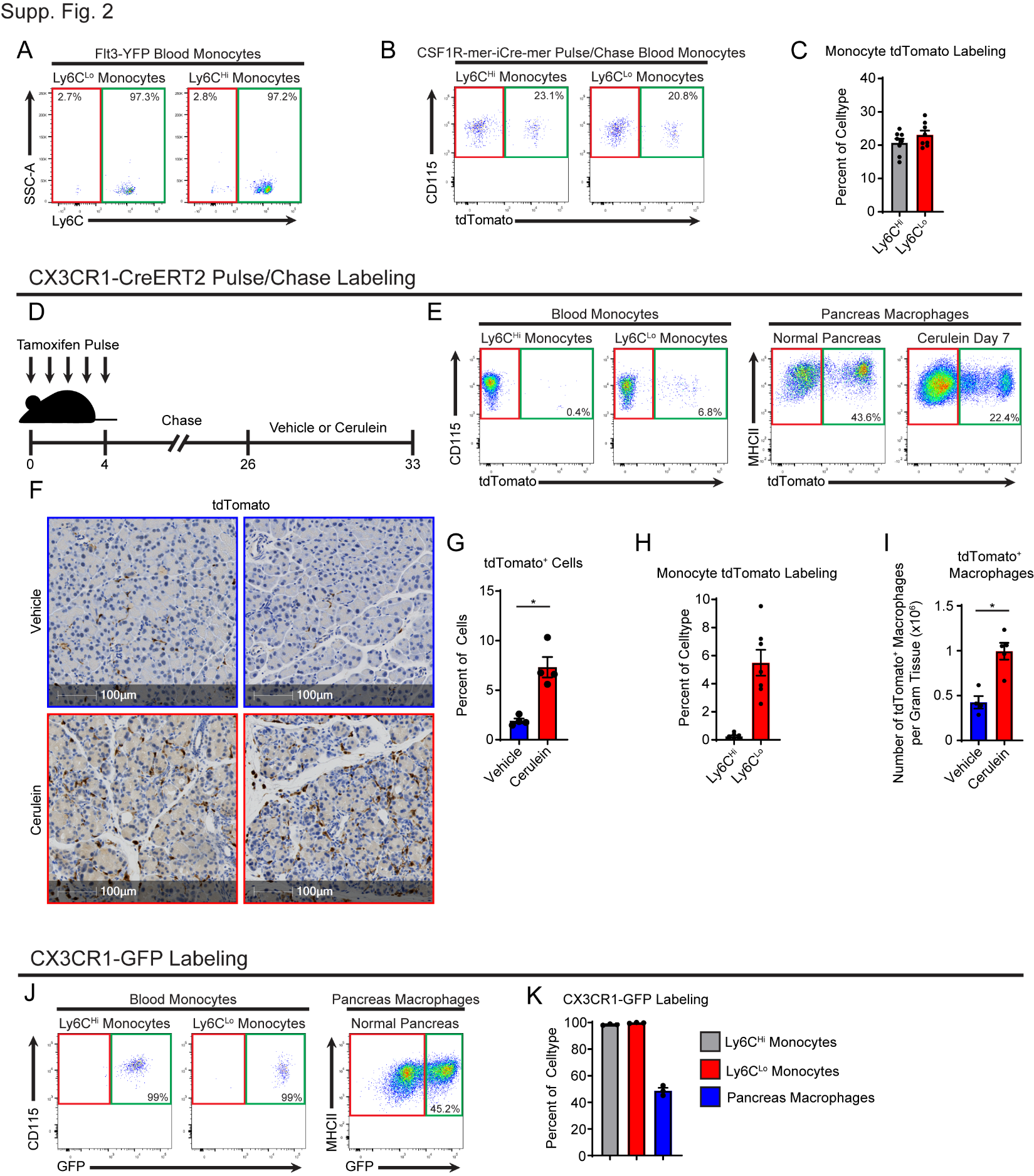

**Supplemental Figure 2: A.** Flow cytometry plots of Flt3-YFP recombination in Ly6C^Lo^ and Ly6C^Hi^ monocytes. **B.** Flow cytometry plots of CSF1R-mer-iCre-mer;LSL-tdTomato recombination in Ly6C^Lo^ and Ly6C^Hi^ monocytes. **C.** Quantification of percentage of Ly6C^Lo^ and Ly6C^Hi^ monocytes expressing tdTomato following tamoxifen pulse/chase. **D.** Schematic of tamoxifen pulse/chase strategy in CX3CR1-CreERT2;LSL-tdTomato. **E.** Flow cytometry plots of tdTomato expression in Ly6C^Lo^ and Ly6C^Hi^ blood monocytes and vehicle and cerulein treated pancreas macrophages. **F.** IHC staining of tdTomato in pancreas tissue of vehicle and cerulein treated CX3CR1-CreERT2;LSL-tdTomato mice pulse/chased with tamoxifen. **G.** Quantification of tdTomato+ cells from (F). **H.** Quantification of percentage of Ly6C^Lo^ and Ly6C^Hi^ monocytes expressing tdTomato. **I.** Density of pancreas macrophages expressing tdTomato in vehicle and cerulein treated mice. **J.** Flow cytometry plots of GFP expression in Ly6C^Lo^ and Ly6C^Hi^ blood monocytes and pancreas macrophages from CX3CR1-GFP mice. **K.** Quantification of percentage of Ly6C^Hi^ and Ly6C^Lo^ blood monocytes and pancreas macrophages expressing GFP.

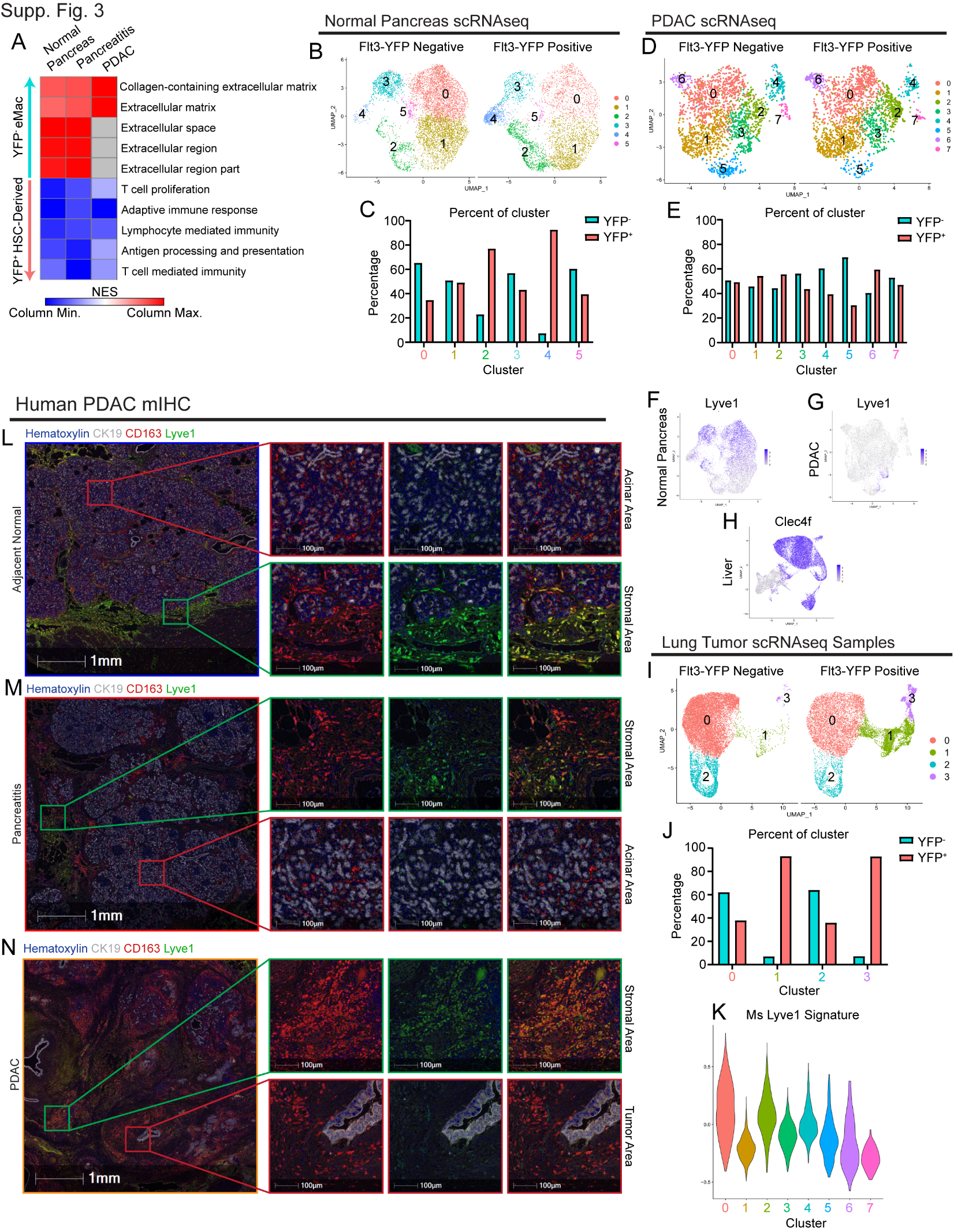

**Supplemental Figure 3: A.** Heatmap displaying normalized enrichment score of significantly enriched gene sets comparing all Flt3-YFP positive versus Flt3-YFP negative macrophages in indicated single-cell data sets, pathways selected by FDR < 0.05. **B.** UMAP plot of scRNAseq analysis of macrophages sorted from homeostatic (normal) pancreas. **C.** Quantification of Flt3-YFP lineage by cluster from UMAP in B, displayed as percentage of each cluster. **D.** UMAP plot of scRNAseq analysis of macrophages sorted from orthotopic KP1 PDAC tumors. **E.** Quantification of Flt3-YFP lineage by cluster from UMAP in D, displayed as percentage of each cluster. **F.** UMAP plot displaying *Lyve1* expression in macrophages from homeostatic (normal) pancreas. **G.** UMAP plot displaying *Lyve1* expression in macrophages sorted from PDAC. **H.** UMAP plot displaying *Clec4f* expression in macrophages sorted from tumor-bearing livers. **I.** UMAP plot of scRNAseq analysis of macrophages sorted from orthotopic KPL lung tumors. **J.** Quantification of Flt3-YFP lineage by cluster from UMAP in F, displayed as percentage of each cluster. **K.** Violin plot showing expression of mouse Lyve1^Hi^ macrophage scRNAseq signature (using top 100 DEGs) mapped into human PDAC macrophage/monocyte data set. **L.** Representative multiplex IHC images from adjacent normal human pancreas samples stained for hematoxylin, CK19, CD163, and Lyve1. Highlighted areas of acinar or stromal area. **M.** Representative multiplex IHC images from pancreatitis samples stained for hematoxylin, CK19, CD163, and Lyve1. Highlighted areas of acinar or stromal area. **N.** Representative multiplex IHC images from PDAC tumor samples stained for hematoxylin, CK19, CD163, and Lyve1. Highlighted areas of acinar or stromal area.

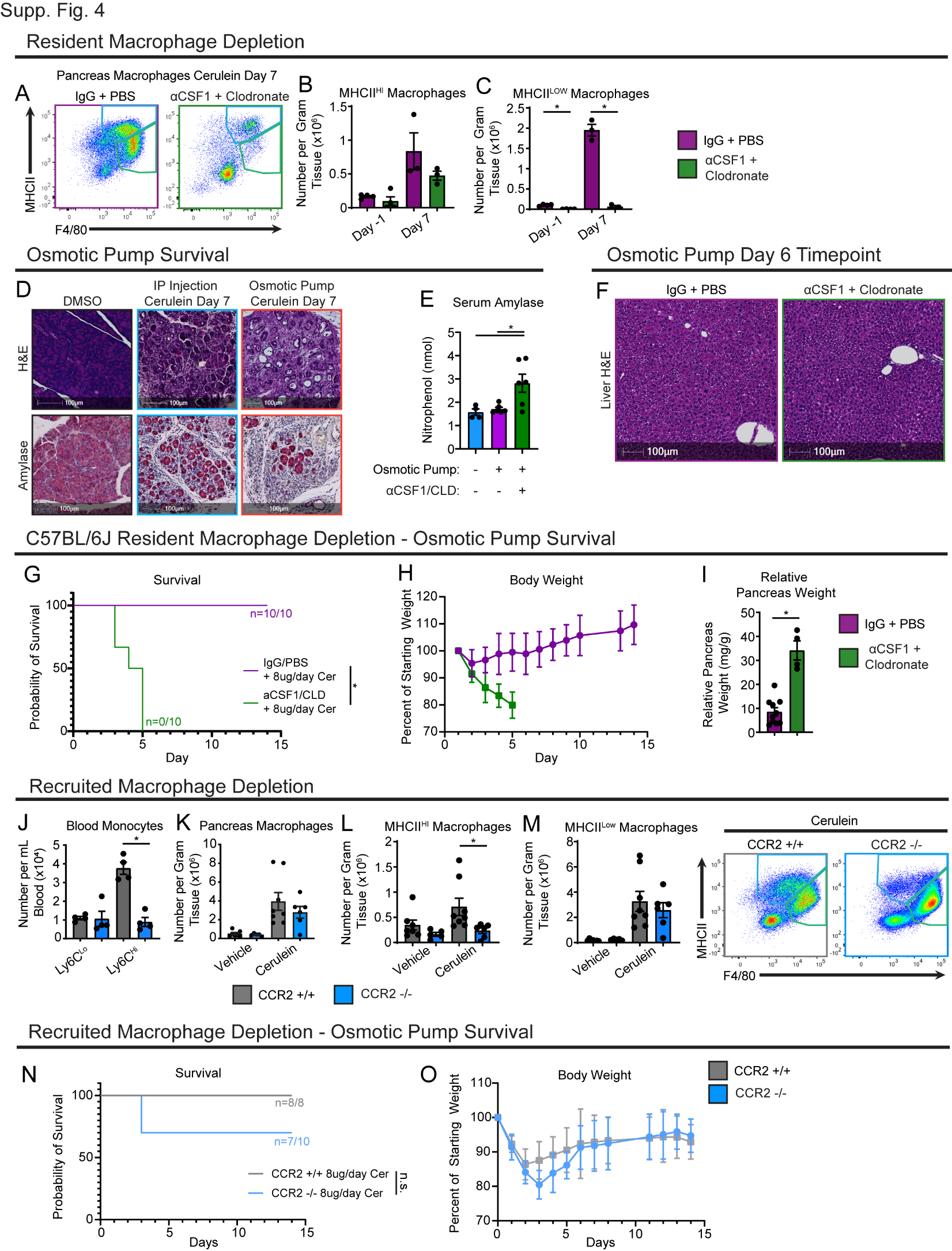

**Supplemental Figure 4: A.** Flow cytometry plot of pancreas macrophages in IgG-PBS or αCSF1-Clodronate mice treated with cerulein for 7 days. **B-C.** Density of MHCII^Hi^ (B) and MHCII^Lo^ (C) macrophages from pancreas of IgG-PBS or αCSF1-Clodronate mice either one day before cerulein start (Day -1) or treated with cerulein for 7 days (Day 7). **D.** Representative IHC images of H&E and amylase stain in pancreas from mice implanted with DMSO loaded osmotic pump, cerulein treated by intraperitoneal injection, and cerulein loaded osmotic pump. **E.** Bar graph showing serum amylase level in mice implanted with or without cerulein loaded osmotic pumps and treated with or without αCSF1-Clodronate. **F.** Representative IHC images of H&E stained liver tissue from IgG-PBS and αCSF1-Clodronate treated mice implanted with cerulein loaded osmotic pumps. **G.** Kaplan-Meier survival curve showing IgG-PBS and αCSF1-CLD groups in C57BL/6 background implanted with cerulein loaded osmotic pumps. **H.** Mouse body weight measurement following implantation of osmotic pump for cerulein delivery, with and without αCSF1-CLD depletion. **I.** Bar graph of relative pancreas weight in IgG-PBS and αCSF1-CLD groups in C57BL/6 background implanted with cerulein loaded osmotic pumps. **J.** Quantification of blood Ly6C^Lo^ and Ly6C^Hi^ monocytes in CCR2+/+ and CCR2-/- mice. **K-M.** Density of total (K), MHCII^Hi^ (L), and MHCII^Lo^ (M) pancreas macrophages in vehicle and cerulein treated CCR2+/+ and CCR2-/- mice. **N.** Kaplan-Meier survival curve showing CCR2+/+ and CCR2-/- mice implanted with cerulein loaded osmotic pumps. **O.** Mouse body weight measurement following implantation of cerulein loaded osmotic pumps.

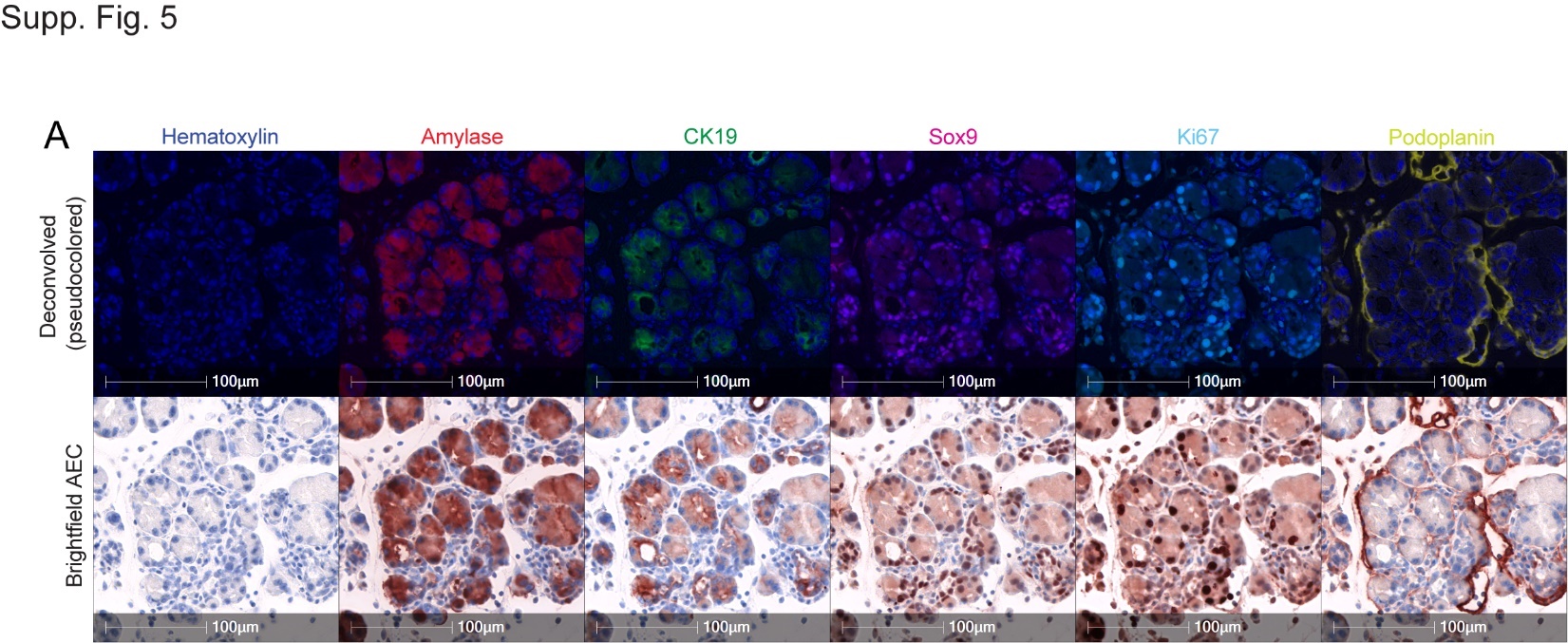

**Supplemental Figure 5: A.** Representative multiplex IHC pseudo-colored (top) or brightfield (bottom) images from pancreas tissue from cerulein treated mice stained for hematoxylin, amylase, CK19, Sox9, Ki67, and Podoplanin.

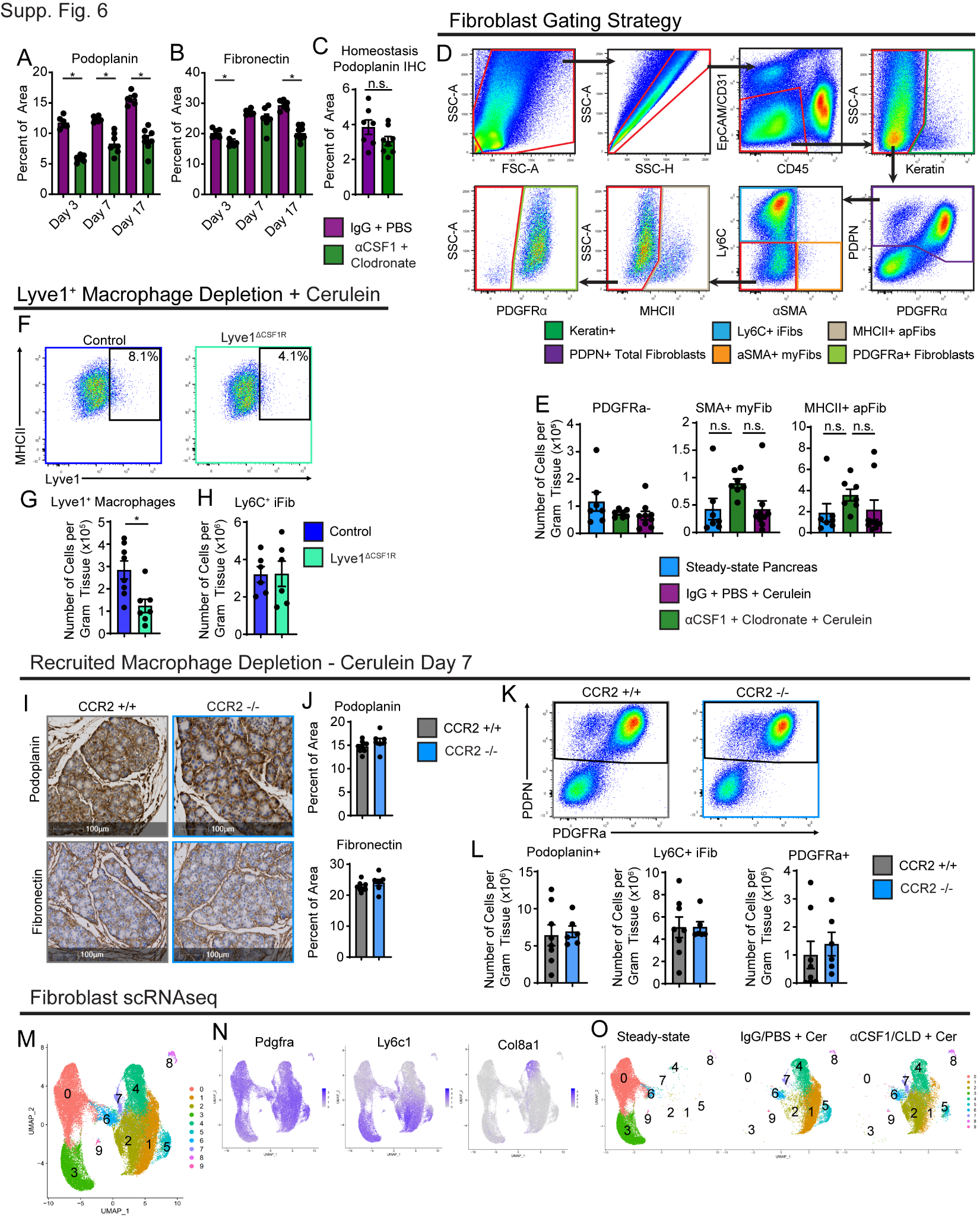

**Supplemental Figure 6: A-B.** Quantification of podoplanin (A) and fibronectin (B) IHC stains on IgG-PBS or αCSF1-Clodronate mice treated with cerulein for 3, 7, or 17 days. **C.** Quantification of podoplanin IHC stain on pancreas tissue of mice treated with IgG-PBS or αCSF1-Clodronate following 10-day recovery period, without cerulein treatment. **D.** Gating strategy for fibroblasts and fibroblast subsets. **E.** Density of PDGFRα(-), α-SMA+, and MHCII+ fibroblasts in pancreas of homeostatic, or IgG-PBS and αCSF1-Clodronate mice treated with cerulein for 7 days. **F.** Flow cytometry plots of Lyve1^+^ macrophages in pancreas tissue of littermate controls, or Lyve1^ΔCSF1R^ mice treated with cerulein for 7 days. **G.** Density of Lyve1^+^ pancreas macrophages in littermate controls, or Lyve1^ΔCSF1R^ mice treated with cerulein for 7 days. **H.** Density of Ly6C^+^ iFibs in pancreas of littermate controls, or Lyve1^ΔCSF1R^ mice treated with cerulein for 7 days. **I.** Representative IHC images of podoplanin and fibronectin stain on pancreas tissue of CCR2+/+ and CCR2-/- mice treated with cerulein for 7 days. **J.** Quantification of (I) podoplanin and fibronectin stains. **K.** Flow cytometry plots of total fibroblasts in pancreas tissue of CCR2+/+ and CCR2-/- mice treated with cerulein for 7 days. **L.** Density of total fibroblasts, Ly6C^+^ iFibs, and PDGFRα^+^ fibroblasts in pancreas tissue of CCR2+/+ and CCR2-/- mice treated with cerulein for 7 days. **M.** UMAP plot of fibroblasts from steady-state pancreas, IgG-PBS and αCSF1-Clodronate mice treated with cerulein for 7 days. **N.** Gene expression of fibroblast marker genes *Pdgfra*, *Ly6c1*, and *Col8a1*. **O.** UMAP plot split by sample (steady-state, IgG-PBS, and αCSF1-CLD).

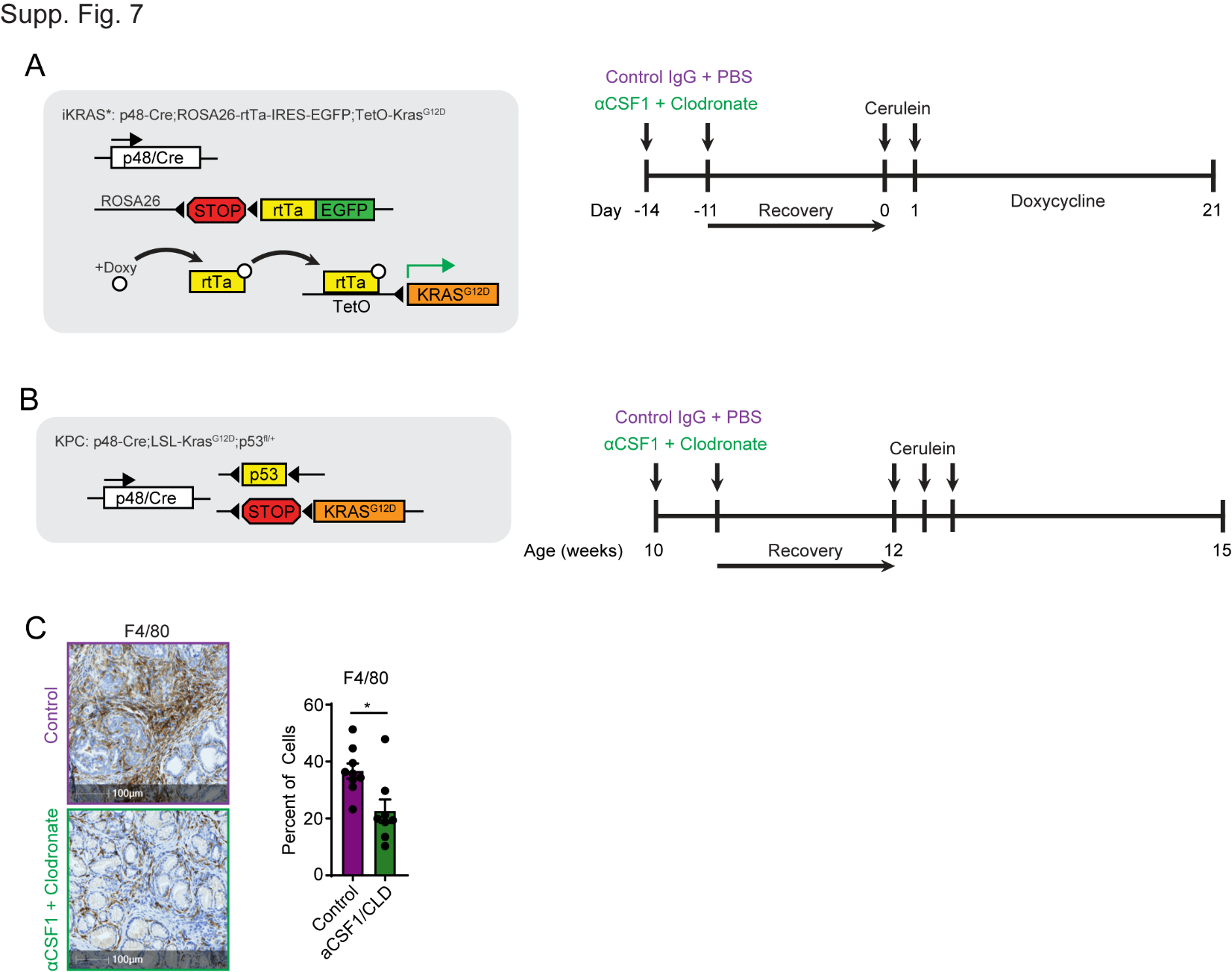

**Supplemental Figure 7: A.** Genetic loci for iKRAS* model and macrophage depletion and doxycycline administration scheme. **B.** Genetic loci of KPC model and macrophage depletion scheme. **C.** Representative images and quantification of F4/80 IHC staining on control and αCSF1-Clodronate treated KPC mice.

Supplementary Tables

Supplementary Table 1: Single-cell RNA-sequencing libraries and cell counts

| Mouse Genotype/Sample | Pathologic Condition | Number of Cells |
| --- | --- | --- |
| Flt3-YFP^+^ Macrophages | Homeostatic pancreas | 4545 |
| Flt3-YFP^(-)^ Macrophages | Homeostatic pancreas | 6868 |
| Flt3-YFP^+^ Macrophages | Pancreatitis (cerulein) | 3594 |
| Flt3-YFP^(-)^ Macrophages | Pancreatitis (cerulein) | 4175 |
| Flt3-YFP^+^ Macrophages | Orthotopic PDAC Tumor | 2297 |
| Flt3-YFP^(-)^ Macrophages | Orthotopic PDAC Tumor | 7213 |
| Flt3-YFP^+^ Macrophages | PDAC metastasis to liver | 5990 |
| Flt3-YFP^(-)^ Macrophages | PDAC metastasis to liver | 13905 |
| Flt3-YFP^+^ Macrophages | Orthotopic Lung Tumor | 11972 |
| Flt3-YFP^(-)^ Macrophages | Orthotopic Lung Tumor | 14605 |
| FvBn Fibroblasts | Homeostatic pancreas | 17880 |
| FvBn Fibroblasts | Pancreatitis + IgG-PBS | 14502 |
| FvBn Fibroblasts | Pancreatitis + αCSF1-CLD | 13145 |
